# Transgenic dCas9-DNMT3A mouse lines enable *ex vivo* and *in vivo* methylation editing with locus-dependent transcriptional effects

**DOI:** 10.64898/2026.02.26.708277

**Authors:** Maria Kalomoiri, Chiara Sorini, Sebastiaan Vos, Anderson Camargo, Alesia Tsalikou, Aleksa Krstic, Chandana Rao Prakash, Per Svenningsson, Maria Needhamsen, Majid Pahlevan Kakhki, Lara Kular, Maja Jagodic

## Abstract

The number of epigenome-wide association studies linking CpG DNA methylation with disease, traits and exposures, continues to rise. Despite the rapid development of epigenome editing tools, establishing causation remains challenging, particularly *in vivo*. In this study, we developed and characterized three Cre-dependent CRISPR-based mouse lines that enable locus-specific DNA methylation deposition by either constitutive or inducible dCas9-DNMT3A expression. We demonstrate robust highly locus-specific DNA methylation deposition at MHC class II (*H2-Ab1)* and interleukin 6 (*Il6)* genes in bone marrow-derived myeloid cells *ex vivo*. Moreover, neuron-specific methylation targeting resulted in reduced cannabinoid receptor 1 (*Cnr1*) expression in striatal neurons *in vivo*. Notably, we demonstrate that the causal effect of DNA methylation on gene expression is locus-dependent, reinforcing the necessity of such editing tools for detailed understanding of the role of DNA methylation and for addressing the causality of disease-associated CpGs.

## Introduction

DNA methylation is a major epigenetic modification involved in the regulation of gene expression, genome integrity, cellular identity, and tissue homeostasis. Large-scale epigenome-wide association studies (EWAS) have identified thousands of differentially methylated CpGs associated with immune, metabolic, neuropsychiatric, and neurodegenerative diseases ^1–5^. However, despite the increasing number of disease-associated methylation signatures, establishing whether specific DNA methylation changes exert causal regulatory effects remains challenging. Most of the studies rely on correlations on bulk tissue, while the role of locus-specific DNA methylation through experimental approaches remains elusive, specifically in *in vivo* assessments.

The emergence of CRISPR-based epigenome editing technologies has enabled targeted manipulation of DNA methylation without altering the underlying DNA sequence. Fusion of catalytically inactive Cas9 (dCas9) to the catalytic domain of DNA methyltransferases, such as DNMT3A, allows methylation deposition at defined genomic loci through guide RNA (gRNA)-directed targeting ^6–13^. Such approaches have provided valuable insights into the regulatory potential of specific CpGs and promoters in cultured cells and more recently *in vivo* following viral delivery strategies^2,13–21^. Nevertheless, efficient implementation of targeted methylation editing in mammalian tissues remains technically challenging due to the large size of epigenome editing constructs, variability in viral delivery, tissue heterogeneity, and uncertainty regarding the stability and functional consequences of induced methylation changes^22,23^.

While the use of dCas9-Dnmt3a transgenic mouse line for targeting of highly methylation-sensitive locus in the liver has alleviated some of these challenges^24^, the generalizability across tissues, cell types (due to mosaic transgene expression), as well as genomic contexts remains incompletely understood. Increasing evidence suggests that the transcriptional effects of DNA methylation are highly context dependent and influenced by chromatin accessibility, regulatory architecture, transcription factor occupancy, and cellular state ^25,26^. Indeed, methylation deposition at selected CpGs does not universally result in transcriptional repression ^27^, underscoring the need for experimental systems capable of functionally interrogating human disease-associated methylation changes directly *in vivo* and in primary cells.

To address these limitations, we generated and characterized novel Cre-dependent transgenic mouse lines enabling constitutive or inducible expression of a dCas9-DNMT3A fusion protein for locus-specific DNA methylation editing. By combining these models with viral delivery of gRNAs, we investigated targeted methylation deposition across several loci, tissues, and experimental settings both *ex vivo* and *in vivo*. We demonstrate that targeted methylation editing can be achieved in immune and neuronal contexts, while the transcriptional consequences of induced methylation remain strongly locus- and context-dependent. Collectively, these mouse models provide versatile platforms for functional interrogation of DNA methylation *in vivo* while highlighting important biological and technical constraints associated with epigenome editing approaches.

## Results

### Generation of the mouse lines and induction of *dCas9-DNMT3A* expression

To generate an inducible dCas9-DNMT3A mouse line, we first evaluated two Cre-dependent constructs expressing dCas9 fused to the catalytic domain of DNMT3A together with an EGFP reporter under the control of a CAG promoter. Both constructs were sharing core elements including the loxP-flanked STOP sites, 3X FLAG tag, the nuclear localization signals (NLS) flanking dCas9, the codon-optimized catalytic domain of the human DNMT3A, and EGFP sequences, differing only in the use of either an IRES or T2A linker for bicistronic expression (**Suppl. Fig. 1a**). We compared locus-specific deposition of methylation and EGFP expression, by co-transfecting HEK293T cells with each dCas9 variant (IRES or T2A) together with a Cre-expressing plasmid and a plasmid carrying the *BACH2*-targeting gRNA (gRNA 8) expressing mCherry reporter (**Suppl. Fig. 1a, b**). We targeted 13 CpGs in the promoter/exon 1 region of the *BACH2* locus (**Suppl. Fig. 1b**). Similar to previous reports, we observed superior EGFP expression and the expected methylation pattern of the deposited methylation in the *BACH2* locus with dCas9-DNMT3A-T2A-EGFP construct (**Suppl. Fig. 1b, c**) ^10^. Based on these results, the T2A-containing cassette was selected for generation of the transgenic mouse line. The Cre-inducible construct was inserted into the *ROSA26* locus using the CRISPR/Cas mediated zygote microinjection (**Fig. 1a**). In the absence of Cre, we observed minimal *dCas9-DNMT3A* expression across tissues, with expected higher expression in homozygous (dCas9-3A +/+) compared to heterozygous (dCas9-3A +/-) animals (**Fig.1b**).

**Fig. 1.**
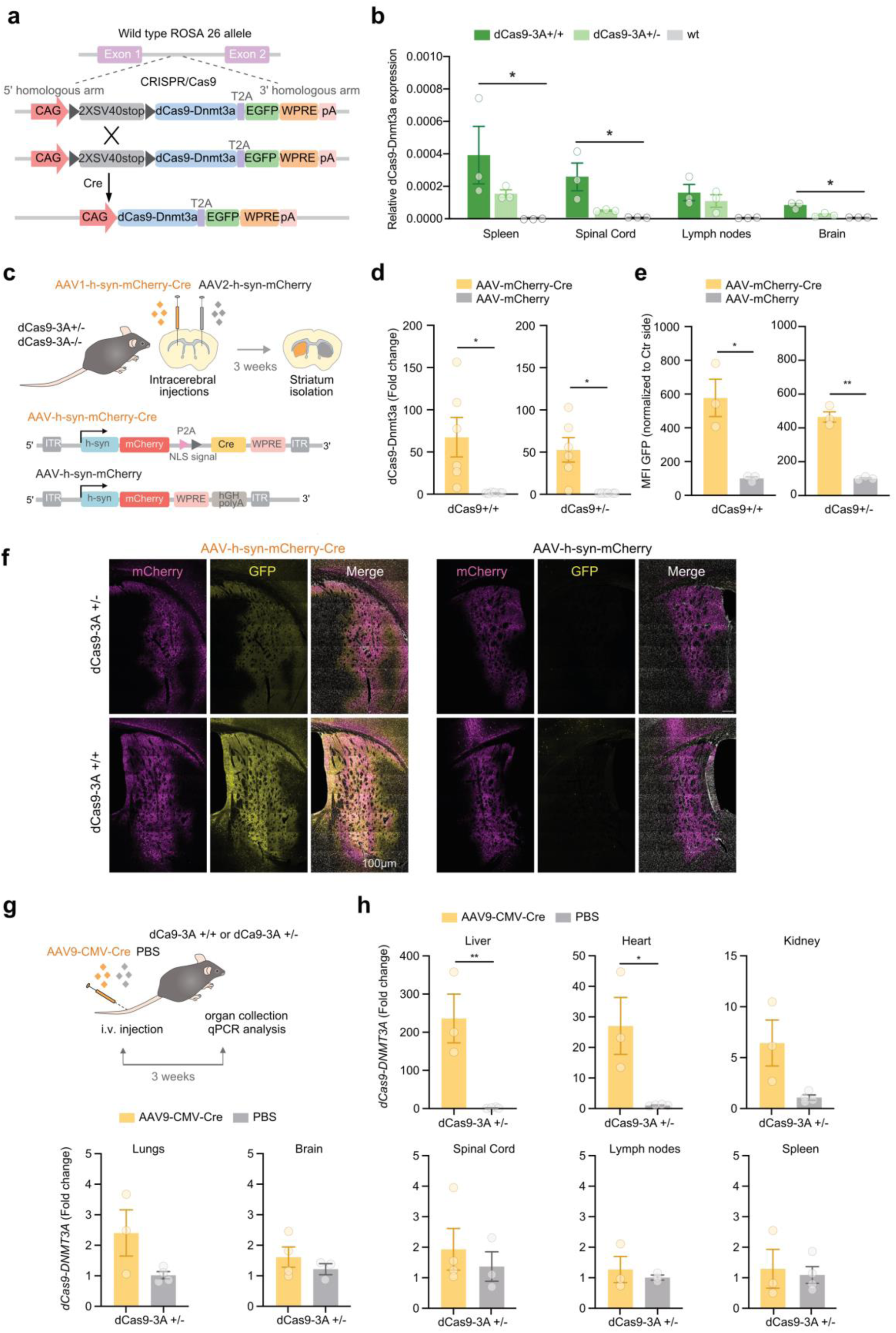
Generation and characterization of dCas9-DNMT3A mouse line. **a**. Schematics of the transgene cassette inserted in the *Rosa26* locus under a CAG-promoter followed by two floxed stop codons, deactivated Cas9 (dCas9), catalytic domain of the human DNA methyltransferase 3 alpha (DNMT3A), the enhanced green fluorescent protein (EGFP) reporter gene, Woodchuck Hepatitis Virus Posttranscriptional Regulatory Element (WPRE) and autocleavage peptide T2A. **b**. Relative to Gapdh *dCas9-DNMT3A* expression assessed by qPCR in different tissues from naïve homozygous (dCas9-3A +/+) and heterozygous (dCas9-3A +/-) dCas9-DNMT3A mice and wild-type (wt) littermate controls. **c.** Schematics of the experimental design of the intracerebral injection of AAV-h-syn-mCherry-Cre virus in the right striatum and the control AAV-h-syn-mCherry virus in the left striatum. The animals were sacrificed 3 weeks after injections and tissues were isolated for qPCR or immunofluorescence. Relative *dCas9-DNMT3A* expression (**d**) and quantification of EGFP (**e**) in the striatum of homozygous and heterozygous dCas9-DNMT3A injected animals. **f**. Representative immunofluorescence staining against EGFP (yellow) and mCherry (magenta) from (**e**); scale bar:100 μm. **g**. Schematics of the experimental design of intravenous injections with AAV9-CMV-Cre viruses or PBS control. **h**. Fold change of *dCas9-DNMT3A* expression (normalized to Hprt) assessed by qPCR in the organs harvested from heterozygous dCas9-DNMT3A animals injected with AAV9 or PBS. Data are represented as mean ± S.E.M. with n=3 per group (**b, d, e**). Non-parametric Kruskal Wallis test with Dunn’s correction for multiple comparisons was used for the assessment of *dCas9-DNMT3A* expression (**b**), while paired (**d, e**) and unpaired two-tailed (**h**) t-tests were performed in other analysis (n=3 in the AAV9 group in all organs, and n=4 for PBS controls, except lymph nodes, Brain, Spinal Cord and heart that n=3). P-values are denoted: *p<0.05, **p<0.01. Non-significant values are not noted.

We next tested the inducible expression of the cassette by intracerebral injections of AAV1-h-syn-Cre-mCherry in the right hemisphere striatum, with the control contralateral striatum receiving AAV2-h-syn-mCherry (**Fig. 1c**). Both AAV1 and AAV2 capsids were selected for their potency in infecting central nervous system (CNS) cells and restricted expression to neurons via the h-syn promoter ^28,29^. Bulk qPCR analysis of harvested striata showed that the AAV-Cre-mCherry delivery led to an average 67- and 53-fold increased expression of the *dCas9-DNMT3A* transcripts three weeks after delivery in homozygous and heterozygous animals, respectively, in comparison to the contralateral side (**Fig. 1d**). Successful viral transduction in both striata could be confirmed using immunofluorescence against EGFP with prominent bilateral mCherry signal and increased EGFP in the ipsilateral versus contralateral control side, indicating potent dCas9-DNMT3A protein expression upon Cre induction locally in the CNS (**Fig. 1e, f)**.

We next addressed systemic induction of the cassette in peripheral organs via intravenous injection of AAV9-CMV-Cre into the tail vein (**Fig. 1g, Suppl. Fig. 2a**), leveraging AAV9’s broad organ tropism and its ability to cross the blood-brain barrier ^30^. Gene expression analysis of *dCas9-DNMT3A* in several organs collected 3-weeks post-injection showed 236- and 27-fold increase in the liver and heart, respectively, of AAV9-CMV-Cre injected heterozygous dCas9-3A+/- animals compared to phosphate buffer solution (PBS)-injected controls (**Fig. 1h**). In contrast, limited induction of *dCas9-DNMT3A* could be observed in other organs including kidney, lung, immune (lymph nodes, spleen) and nervous (brain, spinal cord) tissues (**Fig. 1h, Suppl. Fig 2b**). Similar results were found in the liver and heart of homozygous animals (**Suppl. Fig 2b**).

Collectively, these findings demonstrate that the newly generated Cre-dependent dCas9-DNMT3A mouse line enables efficient local and systemic induction of targeted methylation editing machinery following AAV-mediated Cre delivery.

### Potent and mosaic *dCas9-DNMT3A* expression in constitutive and inducible mouse lines

In order to control the expression of dCas9-DNMT3A, we crossed the dCas9-DNMT3A line with two Cre lines to generate CAG-Cre-ER^TM^-dCas9-DNMT3A (CAG; dCas9-3A) animals, in which expression of the cassette can be induced by tamoxifen injection, and b-Actin-Cre-dCas9-DNMT3A (ACTB; dCas9-3A) animals with constitutive expression of the cassette under the human β-actin promoter (**Fig. 2a, Suppl. Fig. 3a, b)**.

**Fig. 2.**
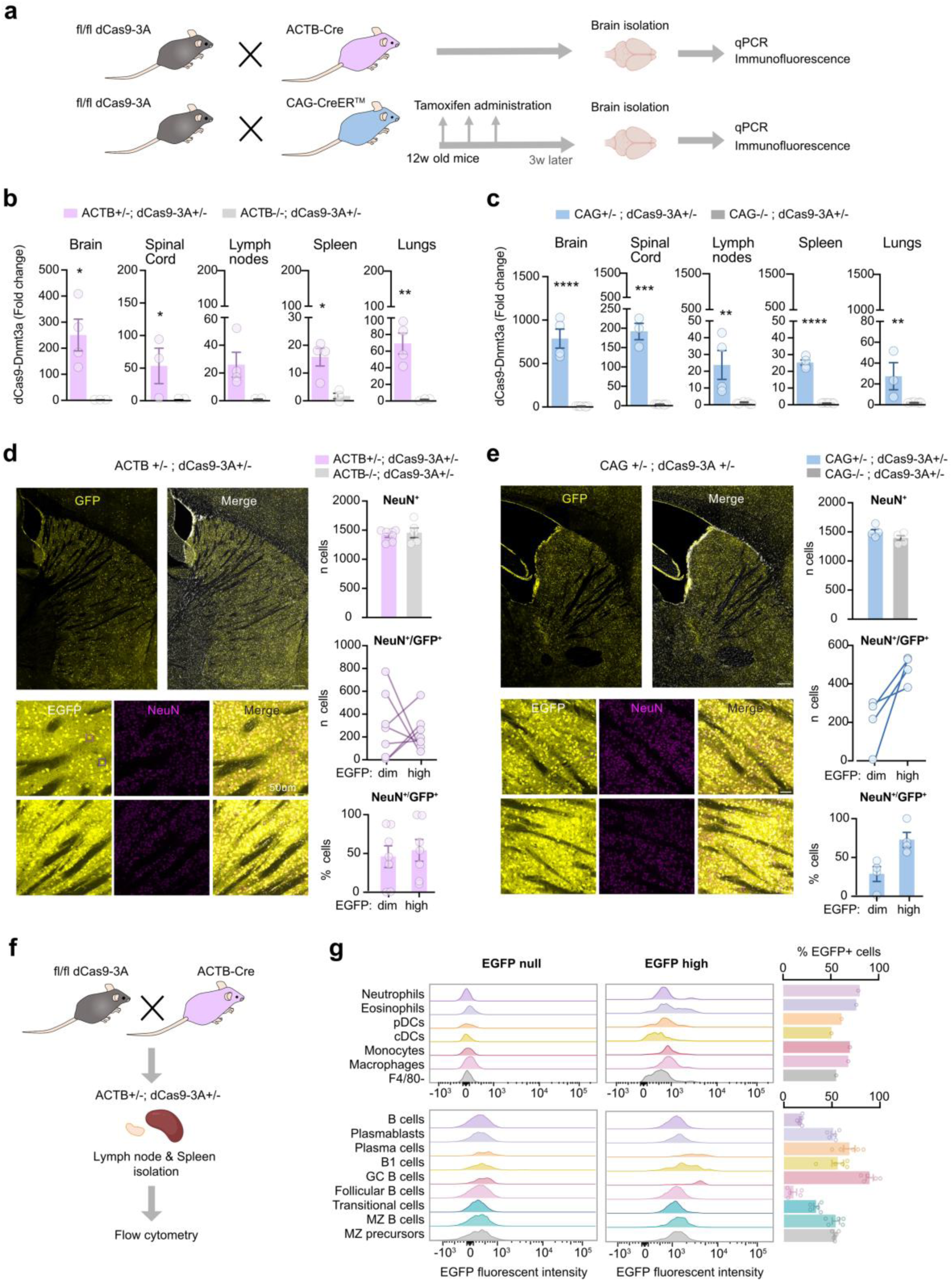
Generation of conditional and inducible dCas9-DNMT3A mouse lines. **a**. Schematic representation of the breeding strategy and the experimental setting for the characterization of the mouse lines that constitutively express *dCas9-DNMT3* under ubiquitous β-actin promoter (ACT; dCas9) or the CAG promoter expression upon tamoxifen treatment (CAG; dCas9). **b**. Relative to Gapdh *dCas9-DNMT3A* expression assessed by qPCR in different organs of ACTB+/-; dCas9-3A +/- (n=3-4 per organ) animals expressed as fold change compared to control ACTB-/-; dCas9-3A +/- (n= 2-3 per organ) animals. **c**. Relative *dCas9-DNMT3A* assessed by qPCR in different organs of CAG+/-; dCas9-3A +/- (n=3-4 per organ) animals expressed as fold change compared to control CAG-/-; dCas9-3A +/- animals (n=7 per organ). Representative of immunofluorescence staining against EGFP in the striatum and quantification of the number of NeuN and/or EGFP positive neurons in the striatum of (**d**) ACTB+/-; dCas9-3A +/- animals (n=7 and 5 animals for ACTB+/- and ACTB-/- groups, respectively), and (**e**) CAG+/-; dCas9-3A +/-animals (n= 4 animals per group), scale bar: 200um (upper), 50um (lower). **f**. Schematics of the experimental design for the immunophenotyping of the lymph nodes and spleen in the ACTB+/-; dCas9-3A +/- animals. **g**. Representative histograms of EGFP expression (null and high) in myeloid and lymphoid cell subpopulations (left) and quantification of the percentage of EGFP-positive cells in 33-weeks old mice (right). Data is represented as Mean ± S.E.M, unpaired t-tests were performed (**b-e**) with P-values marked as *p<0.05, **p<0.01, ***p< 0.001, ****p< 0.0001. Non-significant p-values are not represented.

Phenotypic characterization revealed that ACTB; dCas9-3A mice display breeding difficulties and occasional litter with hydrocephalus, regardless of the genotype. Although body weight at 6 weeks was comparable across genotypes, several animals showed impaired weight gain and reduced size in the adult stage compared to the original dCas9-3A animals which did not exert these features (**Suppl. Fig. 3c-e**). The CAG+/-; dCas9-3A+/- mouse line exhibited reduced litter viability and smaller body size at week 16, but not at week 6, suggesting an impaired weight gain at a later time point (**Suppl. Fig. 3f**). Moreover, tamoxifen administration resulted in occasional mortality.

Characterization of the expression of the cassette in constitutively expressing ACTB+/-; dCas9-3A+/- animals and 3 weeks after tamoxifen administration in inducible CAG+/-; dCas9-3A+/- animals, compared to their respective littermate controls, demonstrated marked but heterogeneous *dCas9-DNMT3A* transcript levels across tissues (**Fig. 2b, c**). In ACTB+/-; dCas9-3A+/- mice, we observed the highest expression levels in the brain (average 250-fold) followed by spinal cord (average 53-fold), lungs, lymph nodes and spleen compared to ACTB-/-; dCas9-3A+/- controls (**Fig. 2b**). On the other hand, CAG+/- ; dCas9-3A+/- mice displayed considerably higher induction of *dCas9-DNMT3A* expression compared to CAG-/-; dCas9-3A +/- controls in brain (787-fold) and spinal cord (200-fold) compared to any other tissue (slightly above 20-fold) (**Fig. 2c**). The cassette expression in the absence of Cre was minimal and comparable among the tissues and lines (**Fig. 2b, c**).

Evaluation of the EGFP expression using immunostaining in the brain, which exhibited the highest expression of *dCas9-DNMT3A*, showed a mosaic EGFP signal in both strains, with cells expressing either bright signal (EGFP-high) or dimmer intensity (EGFP-dim) (**Fig. 2d, e**). A brighter EGFP signal could be found in hippocampal and cerebellar structures of CAG+/-; dCas9-3A+/- compared to ACTB+/-; dCas9-3A+/- mice, consistent with transcript levels (**Suppl. Fig. 4a**). While the total number of NeuN^+^ neurons in the striatum was similar in both transgenic strains, ACTB+/-; dCas9-3A+/- animals displayed a heterogeneous EGFP-expressing neuronal population, with varying EGFP-dim vs. EGFP-high proportions (**Fig. 2d**). On the contrary, tamoxifen-treated CAG+/-; dCas9-3A+/- mice exhibited consistently higher fraction of EGFP-high (72 %) compared to EGFP-dim neurons (28 %) (**Fig. 2e**). Immunostainings of EGFP expression in GFAP^+^ astrocytes, Olig2^+^ oligodendrocytes and Iba-1^+^ microglia revealed no detectable levels of expression in striatum and corpus callosum of both strains (**Suppl. Fig. 4b, c**). This observation could be due to the limited promoter activity in glial cells that would bind in the ACTB-promoter for the induction of the cassette or the insufficient recombination of the cassette.

Similar to the CNS tissue, EGFP levels measured by flow cytometry in the immune compartment varied widely across cell subsets and between animals as exemplified in splenic myeloid and lymphoid cells of ACTB+/-; dCas9-3A strain (**Fig. 2f, g**, gating strategies in **Suppl. Fig. 5, 6**). There were no major differences in frequencies or absolute numbers of myeloid and lymphoid subsets in the lymph node and spleen between the different genotypes of ACTB; dCas9-3A strain (**Suppl. Fig. 7-11**). Likewise, no significant differences in frequencies or absolute numbers of the main myeloid immune cell types were observed in 33-week-old animals across the genotypes, however, the fraction of cells expressing EGFP was lower in old compared to young ACTB+/-; dCas9-3A +/- mice (**Suppl. Fig. 12).** To assess if the animals can mount potent immune response, we immunized them with recombinant myelin oligodendrocyte glycoprotein in adjuvant, used to induce experimental autoimmune encephalomyelitis (EAE) in the original C57BL/6 strain. All mouse lines developed disease with expected clinical features, and no differences were observed across genotypes in young animals (**Suppl. Fig. 13**), while older ACTB+/−; d-Cas9-3A +/− mice displayed more rapid weight loss and the majority of animals had to be sacrificed before the end of experiment (**Suppl. Fig. 13e**).

These data jointly indicate that constitutive and inducible dCas9-DNMT3A mouse lines exhibit robust yet mosaic expression, with interindividual and intercellular variation across CNS and immune tissues. Such heterogeneity likely arises during development and/or recombination and has important implications for downstream functional analyses.

### Genome-wide methylation shows limited off-target deposition of methylation

To further characterize the dCas9-DNMT3A mouse lines, we profiled methylation in spleen and brain tissue from dCas9-3A+/- and wild type (WT) littermate controls of the original strain as well as from tamoxifen-treated CAG+/-; dCas9-3A+/- and CAG-/-; dCas9-3A+/- littermate controls using Illumina arrays that cover over 285,000 CpGs genome-wide. Overall, results showed minimal off-target methyl-deposition and inter-individual variation, as reflected by the overlapping clusters at principal component analyses in each tissue (**Fig. 3a, c**). Importantly, the number and magnitude of differentially methylated CpGs remained low relative to the total number of interrogated sites (>285,000 CpGs), indicating high specificity of the dCas9-DNMT3A system at the genome-wide level.

**Fig. 3.**
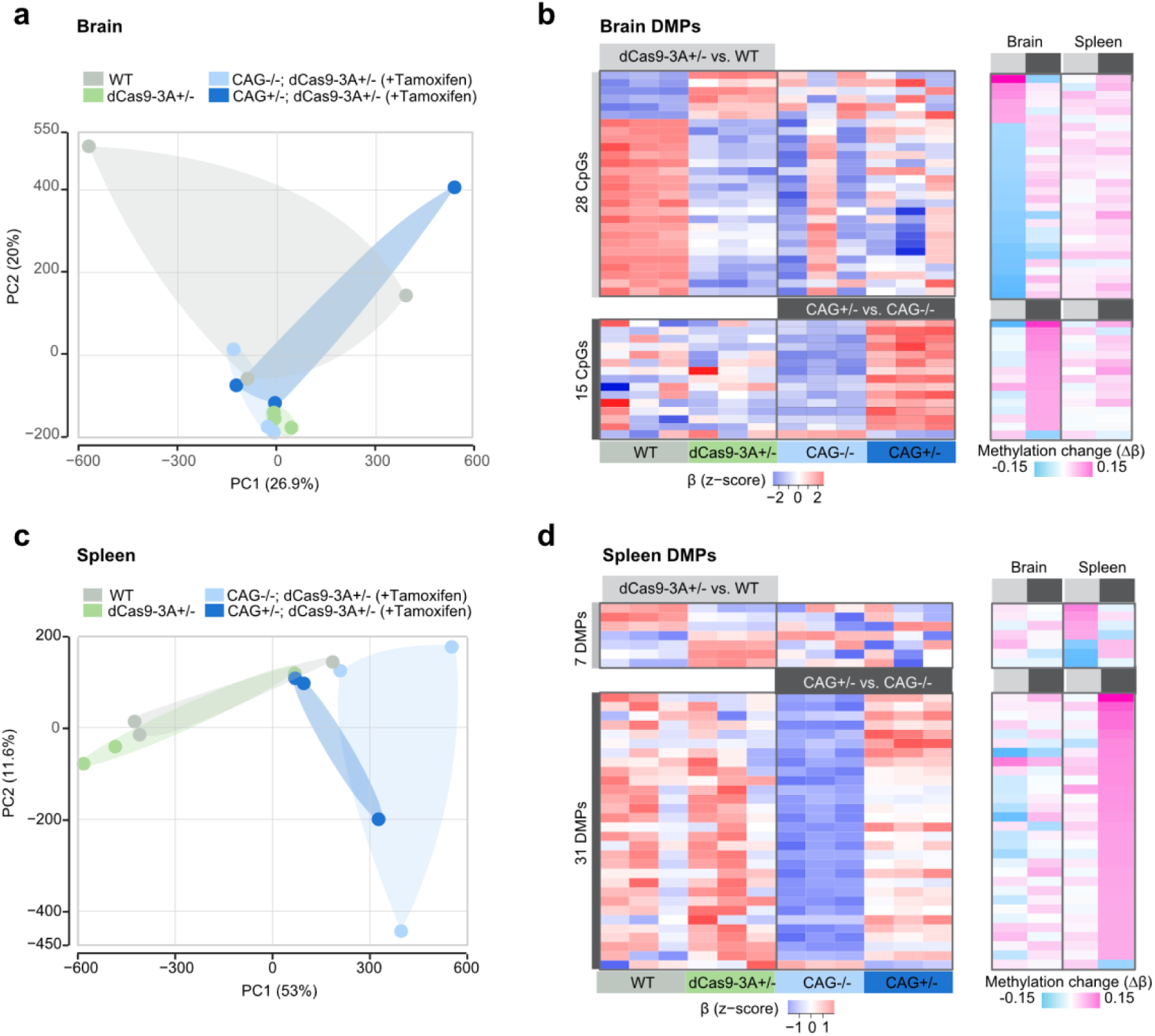
Genome-wide brain and spleen methylome of dCas9-DNMT3A mouse lines. Principal component (PC) analysis of genome-wide methylation levels in brain (**a**) and spleen (**c**) tissues harvested from tamoxifen-treated heterozygous (CAG+/-; dCas9-3A +/-) and control (CAG-/-; dCas9-3A +/-) animals, as well as heterozygous (dCas9-3A +/-) and control wild-type (WT) animals, measured using Illumina Infinium Mouse Methylation BeadChip arrays. A corresponding heatmap of methylation (β) values of differentially methylated CpG positions (DMPs) in CAG+/-; dCas9-3A +/- compared to control CAG-/-; dCas9-3A +/- animals, and dCas9+/- compared to WT animals, in brain (**b**) and spleen (**d**) tissues (*left panel*), and a heatmap of average methylation differences (Δβ) of DMPs across the tissues (*right panel*). Details of array, DMP analysis and outcomes are provided in *Methods* and Supplementary Table 1.

Differential methylation analysis in the brain tissue resulted in 28 and 15 differentially methylated positions (DMPs), displaying methylation differences of > 5% (absolute Δβ > 0.05), between dCas9-3A+/- and WT animals and between tamoxifen-treated CAG+/-; dCas9-3A+/- vs. CAG-/-; dCas9-3A+/- animals, respectively (**Fig. 3b, Suppl. Table 1)**. Most DMPs (78%, 22/28) between dCas9-3A+/- and WT mice were hypomethylated in dCas9+/- animals. In contrast, the majority of DMPs (93%, 14/15) identified in CAG+/-; dCas9-3A+/- compared to CAG-/-; dCas9-3A+/- littermate control mice exhibited hypermethylation, consistent with low off-target methyl-deposition upon activation of dCas9-DNMT3A. These DMPs mapped to TSS of 11 genes including *Adgb*, *Limk2*, *Agmo*, *Olfr165*, *Epas1*, *Gm37013*, *Nek7*, *Unc119b*, *Csmd1*, *Gm5614* and *mir7241* (**Suppl. Table 2**).

Analysis of the spleen yielded 7 and 31 DMPs between dCas9+/- and WT animals and between tamoxifen-treated CAG+/-; dCas9-3A+/- vs. CAG-/-; dCas9-3A+/- animals, respectively (**Fig. 3d, Suppl. Table 1**). The differences between dCas9-3A+/- and WT displayed no preferential directionality of changes and were in intergenic regions. On the contrary and similar to the brain, the vast majority of DMPs (93%, 29/31) in CAG+/-; dCas9-3A+/- versus CAG-/-; dCas9-3A+/- displayed higher methylation following tamoxifen administration. These CpGs affected *Gm47329*, *Rap1gap2*, *Mir3968*, *Trim80*, *Myt1l*, *Pou6f2*, *Gpc5*, *Ly6i*, *Aco2*, *Gm37013*, *Mfsd9*, *Hhat*, *Xirp2*, *Olfr1235*-*ps1*, *Hnf1a*, *Limk1*, *Ints1*, *Lncpint*, *Cdr1os* and *Arhgap6 genes*.

Interestingly, the few changes detected in the brain and spleen were not shared across tissues, suggesting that off-target methylation is both limited and tissue-specific, likely reflecting differences in chromatin accessibility and cellular composition rather than widespread non-specific activity.

### Methylation editing of immune genes in myeloid cells *ex vivo*

We investigated the efficiency of epigenome editing in immune cells *ex vivo* by targeting two immune genes*, H2-Ab1* and *Il6*, in bone marrow-derived macrophages (BMM) after 24h and 6h stimulation with mIFNγ^31^ and LPS (**Suppl. Fig. 14a**), respectively. Macrophages develop as a single adherent monolayer with bone marrow-derived dendritic cells (BMDC) as free-floating cells that are loosely adherent to the macrophage layer (**Suppl. Fig. 14a, b**). As observed *in vivo*, variable levels of *dCas9-DNMT3A* were observed in the cultures of macrophages and dendritic cells, confirming the mosaic nature of expression of the cassette in myeloid cell *ex vivo* (**Suppl. Fig. 14e, f, Suppl. Fig. 15m**).

In these professional antigen-presenting cells, we sought to target the promoter CpGs of the *H2-Ab1* gene encoding MHC class II receptor subunit involved in antigen presentation (**Fig. 4a**). Since *H2-Ab1* promoter did not exhibit a CpG-rich region, we restricted our analysis to the CpGs covering the promoter and part of the exon 1 region by designing two gRNAs (g410 and gTrg2) and pyrosequencing assays jointly covering both target regions in the *H2-Ab1* promoter (**Fig. 4a**). Methylation levels in these *H2-Ab1* promoter CpGs were consistently below 15% in naive BMM and BMDC from original dCas9+/-, constitutive ACTB+/; dCas9-3A+/- and WT animals (**Suppl. Fig. 14g**). We transduced BMM and BMDC from ACTB+/-; dCas9-3A+/- and ACTB-/-; dCas9-3A+/-animals with *H2-Ab1*- or control LacZ-targeting lentiviruses at day 2 of culture and sorted transduced mCherry-expressing cells one week after delivery (**Fig. 3b**, **Suppl. Fig. 14h,i**). Pyrosequencing showed a substantial 3-to-4-fold increase of methylation at CpG positions 3 and 4 of ACTB+/-; dCas9-3A+/- cells transduced with g410 (reaching on average 50 and 60% methylation levels, respectively) compared to gLacZ gRNA, non-transduced cells or ACTB-/-; dCas9-3A+/- cells (**Fig. 4c**). Despite robust methyl-deposition in these 2 CpGs, *H2-Ab1* expression remained unchanged (**Fig. 4d**), implying that methylation at these CpGs is not sufficient to modulate transcription in primary myeloid cells. Transduction with higher viral titers of g410 in antigen presenting cells of both ACTB+/-; dCas9-3A+/- and CAG+/-; dCas9-3A+/- mice resulted in downregulation of *H2-Ab1* gene which was similar to the effect of control gRNA and therefore likely reflected immune response to high viral titer (**Suppl. Fig. 14j,k** and **Suppl. Fig. 15 a-i**). Combined administration of both gRNAs did not allow efficient methyl-deposition at any target regions nor specific gene expression changes (**Suppl. Fig. 15j-m**). Thus, efficient methylation deposition in the targeted CpGs of *H2-Ab1* locus did not associate with the change in gene expression.

**Fig. 4.**
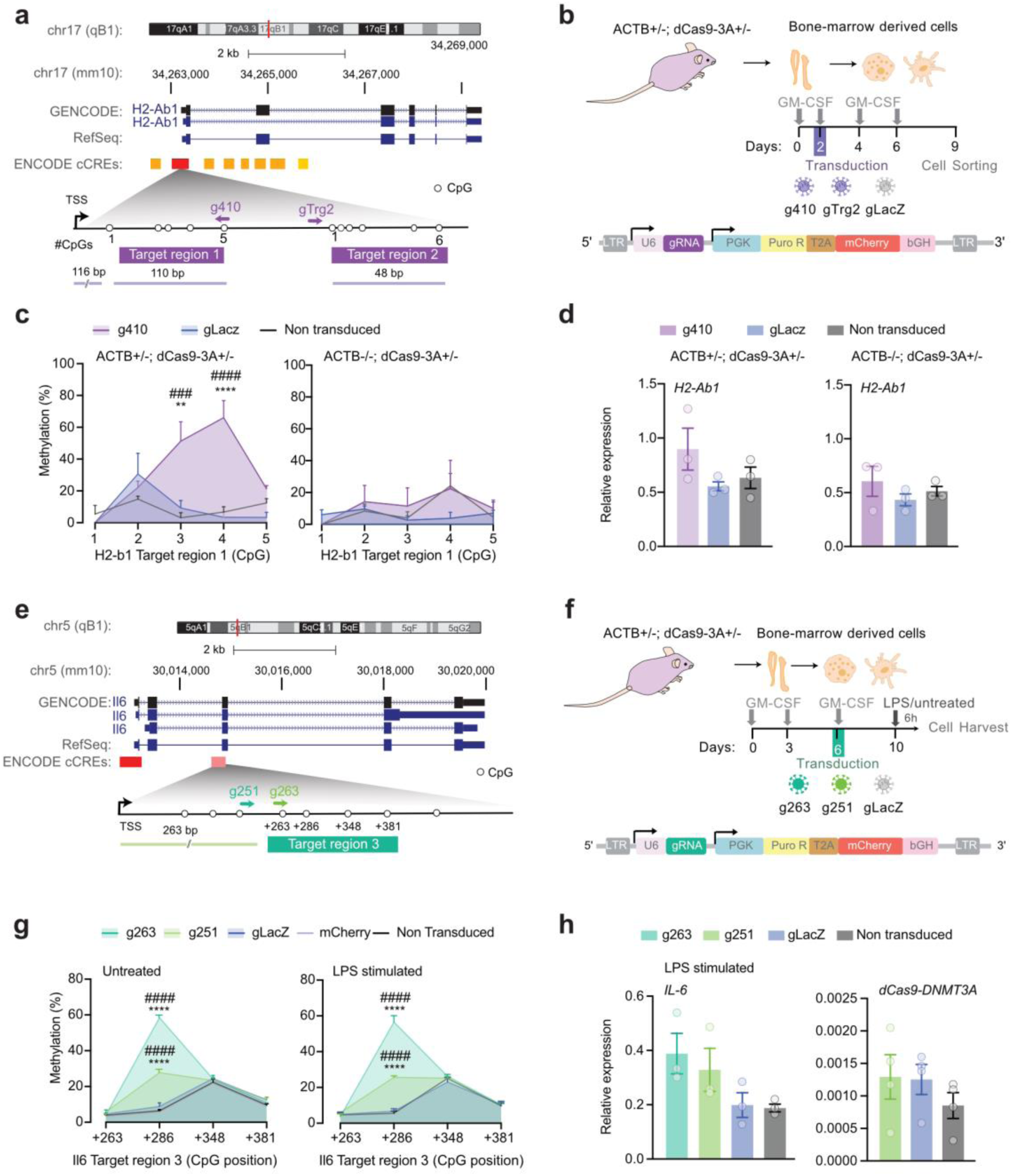
*Ex vivo* targeting of *H2-Ab1* and *Il6* loci in myeloid cells. **a**. Genomic localization of the *H2-Ab1* gene promoter and associated CpGs targeted by g410 and gTrg2 gRNAs. Annotation of regulatory features, *i.e.*, CpG island (CGI) and chromatin state segmentation by hidden Markov model (ChromHMM) from ENCODE/Broad (dark and light red indicates active and poised/weak promoter). **b**. Schematics of the experimental design of lentiviral transduction with targeting g410 or gTrg2 and control LacZ gRNA and sorting of bone marrow-derived cells from the mouse line that constitutively expresses *dCas9-DNMT3A* (ACTB+/-; dCas9-3A +/-) and control animals (ACTB-/-; dCas9-3A +/-). **c**. DNA methylation levels of *H2-Ab1* promoter CpGs quantified using pyrosequencing. **d**. Relative to *Gapdh* expression levels of *H2-Ab1* and *dCas9-DNMT3A*. **e**. Genomic localization of the *Il6* gene regulatory CpGs at exon 2 targeted by g251 and g263 gRNAs. **f**. Schematics of the experimental design of lentiviral transduction with targeting g251 or g263 and control LacZ gRNA and sorting of bone marrow-derived cells from the mouse line that constitutively expresses *dCas9-DNMT3A* (ACTB+/-; dCas9-3A +/-) and control animals (ACTB-/-; dCas9-3A +/-). **g**. DNA methylation levels at *Il6* CpGs quantified using pyrosequencing. **h**. Relative expression levels of *Il6* and *dCas9-DNMT3A*. Data are represented as mean ± S.E.M. with n=3 per group (**c, d, g)** and n=4 per group in **h**. Two-Way ANOVA with Tukey’s multiple comparison post-hoc test was performed in c, g. One-Way ANOVA with Tukey’s multiple comparison post-hoc test was performed in d and h. P-values are denoted: *p.adj <0.05, **p.adj <0.01, ***p.adj <0.001, ****p.adj <0.0001, where * in comparison to gLacZ, and # in comparison to non-transduced. Non-significant p-values are not represented.

We next targeted a particular exonic CpG of the *Il6* gene (+286 in Target region 3), previously shown to regulate *Il6* expression in the murine RAW264.7 cell line and alveolar macrophages ^32^ (**Fig. 4e**). The *Il6* exon 2 region displayed intermediate (lower than 50%) methylation levels following stimulation with LPS (**Suppl. Fig. 16a**). Cells from ACTB+/-; dCas9-3A+/- animals were transduced with two *Il6*-targeting gRNAs, g263 or g251, or control viruses three days prior to bulk harvest (**Fig. 4f**). Robust deposition of methylation was achieved by both gRNAs, particularly g263, at CpG +286 in both untreated and LPS-stimulated cells while the surrounding CpGs were unaffected (**Fig. 4g**). At CpG +286, methylation increased from 8% in the gLacZ control cells to 28% and nearly 60% in the g251 and g263 conditions, respectively. Of note, the level of methyl-deposition was associated with the level of *dCas9-DNMT3A* expression (**Suppl. Fig. 16b-e**). Despite sizeable CpG methylation, *Il6* expression did not significantly differ among the conditions (**Fig. 4h**).

To further explore potential off-target effects, we evaluated sequence-dependent specificity of all gRNAs in silico using the CRISPOR tool. The majority of predicted off-target sites, prioritized based on sequence similarity, were located in genomic regions either lacking CpG sites or containing only sparse CpG density, limiting their potential relevance for DNA methylation analysis. Based on previous findings indicating unwanted methylation by dCas9-DNMT3-based constructs at loci with low-to-medium basal methylation levels ^10^, we further experimentally assessed potential off-target methylation at such unrelated (irrelevant to the target gRNA) genomic regions. Specifically, we analyzed *H2-Ab1* and *Cnr1* promoters in samples transduced with *Il6*-targeting gRNAs and conversely, we profiled *Il6* and *Cnr1* loci after delivery of *H2-Ab1*-targeting gRNA. Pyrosequencing indicates no detectable off-target methyl-deposition at unrelated CpG loci under the experimental conditions tested (**Suppl. Fig. 17**).

Thus, methylation editing in immune cells *ex vivo* demonstrates efficient and highly site-specific deposition of methylation at *H2-Ab1* and *Il6* regulatory CpGs, which however did not affect transcription, providing more insights into the regulation of these loci in primary myeloid cells.

### *In vivo* methylation editing leads to downregulation of *Cnr1* gene expression

We next explored *in vivo* methylation editing by targeting the *Cnr1* gene encoding Cannabinoid Receptor 1 (**Fig. 5a**), which is highly expressed in the dorsolateral striatum of juvenile mice^33^). Pyrosequencing confirmed that *Cnr1* promoter is largely unmethylated and thus amenable to targeted methylation, with baseline methylation levels on average below 5% at two different promoter regions encompassing 15 and 7 CpGs, in bulk dorsolateral striatum tissue across all the original and constitutive dCas9-3A strains (**Suppl. Fig. 18a, b**). We designed AAV2-h-syn viruses expressing simultaneously two gRNAs (g1 and g2) that were previously shown to successfully target *Cnr1* promoter region using SadCas9-VPR system^34^ (**Fig. 5a, b**). The gRNAs were modified in the PAM site to adapt to our SpdCas9, as the original gRNAs were designed for the SadCas9 complex. We could confirm the efficient induction of dCas9-3A upon stereotaxic injection of the viruses by immunofluorescence (**Suppl. Fig. 18c, d**). *Cnr1*-targeting (AAV-h-syn-g1g2-mCherry-Cre) and LacZ control (AAV-h-syn-gLacZ-mCherry) viruses were administered via injection into the right and left dorsolateral striatum, respectively, of dCas9-3A+/+ and WT mice and expression analysis was performed 6-7 weeks after injection.

**Fig. 5.**
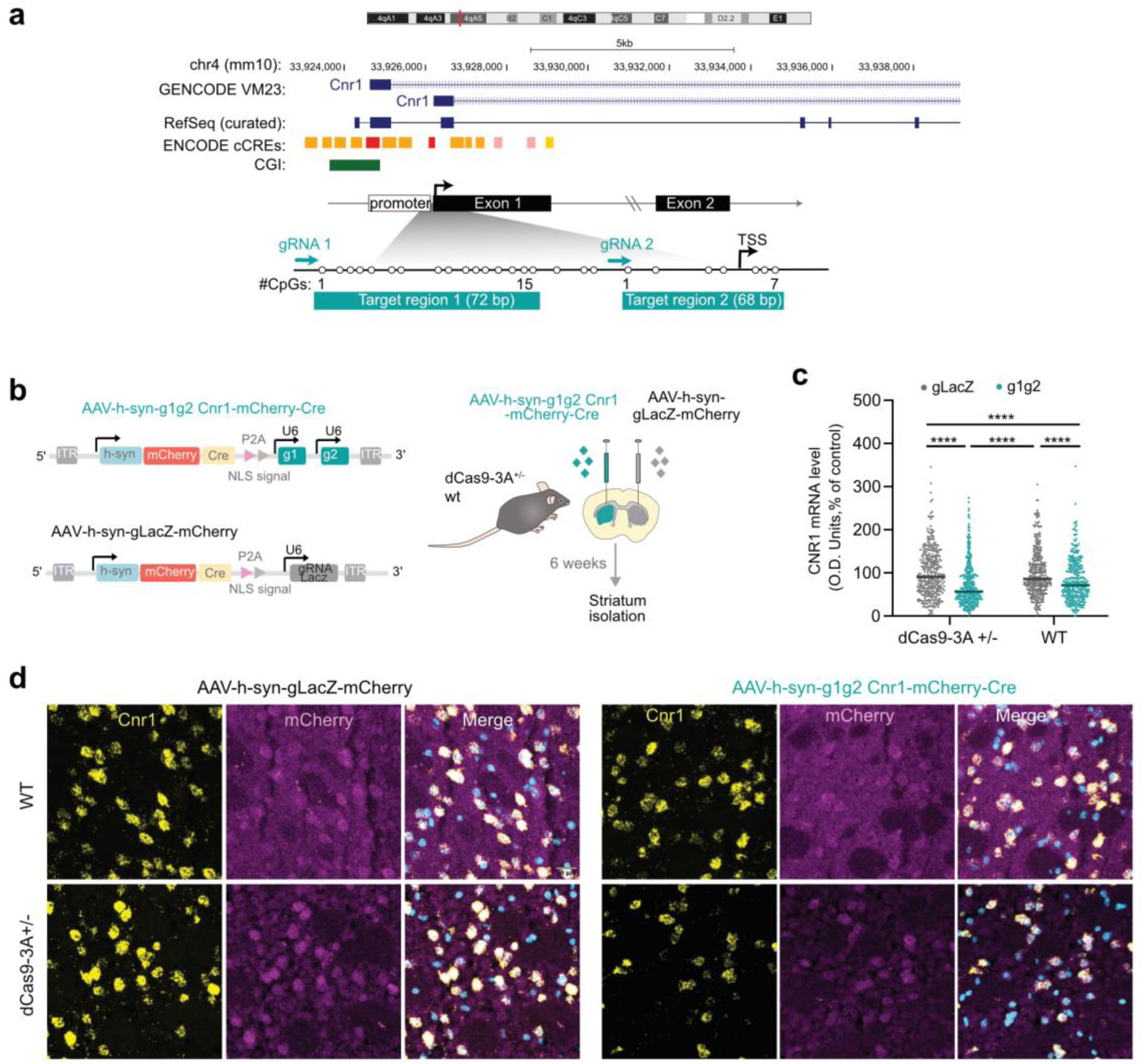
*In vivo* targeting of the *Cnr1* gene in striatal neurons. **a**. Genomic localization of the *Cnr1* gene promoter and associated CpGs targeted by g1 and g2 gRNAs. Annotation of regulatory features, *i.e.*, CpG island (CGI) and chromatin state segmentation by hidden Markov model (ChromHMM) from ENCODE/Broad (red, dark and light orange representing active promoter, strong and weak enhancer, respectively). **b**. Schematic representation of the experimental design of the AAV-hsyn-g1g2-mCherry-Cre injected right hemisphere and the AAV-hsyn-gLacZ-mCherry-Cre injected left hemisphere of the heterozygous dCas9-DNMT3A (dCas9-3A +/-) and wild-type (WT) littermate control animals. **c**. Quantification of *Cnr1* expression using RNAscope (n=350 *Cnr1*-positive cells from 7 animals per group). The median value and individual cell values are represented as bar and dots, respectively. Two-Way ANOVA test with Tukey’s multiple comparison post hoc test was used for comparison, ****p.adj<0.0001. **d**. Representative RNAscope images from the injected animals across the rostrocaudal axis of the striatum. Scale bar: 20 um.

Despite an overall induction of dCas9-DNMT3A cassette, initial bulk analysis of dorsolateral striatum tissue did not show significant difference in *Cnr1* methylation or expression between g1g2 and gLacZ conditions, which likely resulted from heterogenous transduction efficiency observed across the neural cell populations of the striatum (**Suppl. Fig. 18d-g**). We therefore assessed *Cnr1* expression at a single-cell resolution using RNAscope after validation of the method in WT animals (**Suppl. Fig. 18j**). We quantified *Cnr1* expression per cell, 6 weeks after bilateral injections of g1g2- and gLacZ-carrying viruses in the dorsolateral striatum of dCas9-3A +/- and WT control mice, using mCherry expression as a proxy of transduction. Delivery of *Cnr1*-targeting g1g2 led to a significant 25% reduction of the expression of *Cnr1* in striatal neurons of dCas9-3A+/-animals, exceeding the subtle downregulation effect observed in WT animals (**Fig. 5c, d**). These findings indicate that targeted DNA methylation can effectively repress gene expression *in vivo* in a cell-specific manner.

## Discussion

We generated and characterized Cre-dependent dCas9-DNMT3A mouse lines enabling locus-specific DNA methylation editing in both *in vivo* and *ex vivo* settings. Across immune and neuronal systems, targeted methylation deposition was robust and highly specific, while the transcriptional consequences were strongly locus- and context-dependent. These findings highlight both the power and the limitations of epigenome editing approaches for functional interrogation of disease-associated CpGs.

A central advantage of these models is the flexible control of the editing machinery through constitutive or inducible Cre systems, combined with viral delivery of gRNAs. This design circumvents a major limitation of previous approaches requiring co-delivery of large epigenome-editing constructs and enables efficient targeting across tissues. The observed minimal leakage in the absence of Cre further supports the suitability of these lines for controlled experimental manipulation. At the same time, differences in recombination efficiency across tissues upon AAV-based delivery methods underscore the importance of careful experimental design when applying these models in diverse biological contexts. Consistent with the known tropism of AAV9, potent recombination could be observed in the two mostly transduced organs, liver and heart. This tissue bias likely contributes to the prominent use of hepatic targets such as *Pcsk9* in previous *in vivo* and *in vitro* epigenome editing studies ^13,19,20,24,35^, where direct and efficient delivery and editing in liver provide a favorable context for observing robust transcriptional outcomes. These observations highlight that biological conclusions from epigenome editing experiments must be interpreted in light of tissue-specific delivery efficiency and accessibility.

A key strength of the system is its high specificity. Both genome-wide methylation profiling and targeted analyses demonstrate minimal off-target methylation, with changes restricted to intended loci and no detectable gRNA-dependent effects at unrelated CpG sites. This supports the precision of dCas9-DNMT3A-mediated editing *in vivo* and distinguishes it from broader perturbation approaches. The limited and tissue-specific background changes observed are likely attributable to chromatin accessibility and cellular composition rather than non-specific enzymatic activity. In line with our previous findings, off-target methylation does not occur uniformly across the genome but appears to be influenced by intrinsic features of specific loci ^10,36^. In the present study, the absence of substantial off-target effects despite detectable expression of the editing machinery further suggests that genomic context and local accessibility constrain where unintended methylation can occur. Together, these observations indicate that the activity of dCas9-DNMT3A is shaped by both the epigenomic landscape and expression levels, resulting in selective susceptibility of certain loci while leaving the majority of the genome unaffected under physiological conditions.

Our results provide direct experimental evidence that DNA methylation does not exert uniform transcriptional effects across genomic loci. Despite efficient methylation deposition at CpGs within the *H2-Ab1* and *Il6* loci, no consistent changes in gene expression were observed in primary myeloid cells. The downregulation of *H2-Ab1* expression observed following both targeting and non-targeting gRNA delivery is consistent with prior reports describing early transcriptional changes after viral transduction, likely reflecting viral immune evasion ^37^. In contrast, targeting the largely unmethylated promoter of *Cnr1* in neurons resulted in significant transcriptional repression. These findings indicate that the regulatory consequences of DNA methylation depend on local genomic architecture, including CpG position, chromatin accessibility, and transcription factor occupancy ^38^. In particular, methylation at isolated CpGs or within partially methylated regions may be insufficient to impact transcription in general and particularly in complex immune loci such as MHC class II ^10,39^. Together, these observations reinforce the need for locus-specific functional validation of candidate CpGs identified in association studies.

The mosaic expression of the dCas9-DNMT3A cassette represents an important limitation of the current models as both constitutive and inducible strains exhibited substantial intercellular and inter-individual variability ^24,40,41^. This heterogeneity reduces sensitivity in bulk analyses and necessitates single-cell or reporter-based approaches to accurately assess functional outcomes, as illustrated by the requirement for RNAscope to detect *Cnr1* repression in neurons. Additionally, despite the minimal off-target methylation detected at both genome-wide and locus-specific levels, we observed reduced viability and developmental abnormalities in some mouse lines. These findings indicate that epigenome editing outcomes cannot be solely interpreted based on targeting specificity. The presence of an active DNMT3A catalytic domain, even when guided with high precision, may impose a broader epigenetic burden through low-level or stochastic methylation events that remain below detection thresholds in bulk assays. The mosaic expression patterns likely arise from incomplete recombination, variability in transgene regulation such as silencing of the cassette, or cellular selection against high expression levels leading to altered tissue composition and reduced organismal fitness. Importantly, constitutive or leaky expression during development is likely to exacerbate these effects, as DNA methylation plays a central role in lineage specification and tissue homeostasis ^42–45^. Together, these observations underscore that even highly specific epigenome editing systems can have unintended physiological consequences, particularly when expressed broadly or during sensitive developmental windows.

Taken together, these models provide a versatile platform for functional interrogation of DNA methylation in physiologically relevant systems. By enabling targeted manipulation of specific CpGs, they offer a powerful approach to directly test causal relationships. However, our findings also emphasize that the effects of DNA methylation are highly context-dependent and cannot be generalized across loci. Future work integrating systematic gRNA screening, single-cell readouts, and improved control of transgene expression will be essential to fully exploit the potential of epigenome editing for understanding gene regulation and developing therapeutic strategies.

## Methods

### Generation of Cre-dependent dCas9–DNMT3A transgenic mouse lines

The conditional mouse strain dCas9-DNMT3A was generated by Biocytogen on the C57BL/6J genetic background using the CRISPR/Cas9 technology, targeting the insertion of the cassette in the *Rosa26* locus utilizing the gRNA AAGGCCGCACCCTTCTCCGG. The cassette that was used for the generation of mice had the following sequence: a CAG promoter, a loxP site, a SV40 late polyA signal (pA) followed by the signal stop cassette lox2-STOP sequence and another loxP site, a 3X FLAG sequence, an SV40 nucleoporin (NLS), the deactivated Cas9 (dCas9), another NLS and the catalytic domain of the human methyltransferase, followed by a T2A peptide, an EGFP reporter and the woodchuck hepatitis post-transcriptional (WPRE). In short, CAG-SV40-Lox2-STOP-3XFLAG-SV40NLS-dCas9-NLS-DNMT3A-T2-EGFP-WPRE (**Suppl. Fig.1a**). We chose the catalytic domain of the human methyltransferase tethered with the dCas9 for the generation of mice, to mimic the physiological human condition of the human methyltransferase and generate “humanized” mice. We bred the original mice to generate conditional heterozygous (dCas9-3A +/-) and homozygous (dCas9-3A +/+) animals for testing. The original floxed dCas9-3A strain was bred with mice expressing CreER^TM^ (Stock# 004682) ^46^ under tamoxifen administration (named CAG-CreER^TM^; dCas9-DNMT3A^fl/fl^) or mice that were constitutively expressing Cre, utilizing the human β-Actin promoter, ACTB-Cre mice (Stock# 019099), purchased from the Jackson Laboratory. Homozygous and heterozygous animals for the dCas9-DNMT3A construct and heterozygous for the CAG-CreERT or ACTB-Cre expression were generated for experiments. Mice floxed for the dCas9-DNMT3A and carrying no copy of the pCAG-CreERT and ACTB-Cre alleles, were used as control animals in the experiments. All primers used for the PCR genotyping of the three mouse lines are listed in **Suppl. Table 2**.

The animals were kept in rooms with 12-h light/dark cycles and controlled temperature/humidity (20 °C/53%). Food pellets and water were given to animals on an ad libitum basis. The local ethical committee at Karolinska Institutet approved the experiments (9328-2019/2137-2022) and the experiments were conducted in accordance with the European Communities Council Directive of 24 November 1986 (86/609/EEC).

### Tamoxifen-inducible activation of dCas9-DNMT3A

The inducible expression of the dCas9-DNMT3A cassette was accomplished by the administration of tamoxifen in CAG+/-; dCas9-3A +/- or CAG-/-; dCas9-3A +/-. Tamoxifen (Sigma, T5648) was diluted in sunflower corn oil (S5007, Sigma) and ethanol at a ratio of 9:1. The final concentration of tamoxifen was 10mg/ml and it was administered intraperitoneally to animals at 2mg per day for 5 days, every other day. For the *ex vivo* cultures, 4-OH Tamoxifen was used at a concentration of 500 nM in a single dose in CAG+/-; dCas9+/- mice at day 3 of culture.

### Adeno-associated viruses (AAVs)

AAV1-hSyn-mCherry-P2A-Cre-WPRE (#107312) and AAV2-hSyn-mCherry (#114472) viruses were purchased from Addgene. The stock concentration of the viruses was 1.3 X 10^13^ vg/ml and 1.8 X10^13^ vg/ml, respectively. AAV2-g1g2-Cre-mCherry and AAV2-gLacZ-Cre-mCherry were customized and purchased from VectorBuilder and the stock concentration of these viruses was 6.64 x 10^13^ vg/ ml and 6.62 x 10^3^ vg/ml, respectively. The AAV2-g1g2-Cre-mCherry and AAV2-gLacZ-Cre-mCherry were diluted in PBS before administration to the animals at a final concentration of 3 x 10^13^ vg/ml. For the intravenous injections, the CMV-Cre-AAV9 (#105537) virus was used at a single dose of 2 x 10^11^ vg per mouse. The stock concentration of the virus was 2.3 x 10^13^ vg per ml.

### Stereotaxic surgery on mice

Mice that were 3-4 months old were used for intracerebral injections. Prior to the procedure, the animals received subcutaneous analgesia with Temgesic at 1 mg/kg. The mice were anesthetized with 3% isofluorane at the initiation of the process before the operation and at 2% for the maintenance with 0.5 lpm air flow. The mice were restrained in a stereotactic frame with their heads fixed to the stereotactic frame. Occulentum was applied to their eyes to avoid dryness and xylocaine as local skin anesthetic. The coordinates for the intrastriatal injections of the AAV1-hSyn-mCherry-P2A-Cre-WPRE and AAV2-hSyn-mCherry viruses were anterior-posterior (AP): +0.5mm, medial-lateral (ML): +/- 2.0 mm and dorsoventral (DV): -2.5 mm relative to bregma and dural surface, while the coordinates for the injection of the AAV-Cnr1-Cre-g1g2 and AAV-Cre-gLacZ targeting the dorsolateral striatum were AP): +0.5mm, medial-lateral (ML): +/- 2.5 mm and dorso-ventral (DV): -3.5 mm. 10 μl Hamilton syringes were used for the injection. One microliter of viruses was injected to each striatum of the mice with a rate of 0.2 μl/min for 5 min. At the end of the procedure the needle was kept for additional 5 min and then slowly removed. After the operation, the skin was sewed with sutures, and the mice were observed until they woke up. Mice were injected subcutaneously with 0.5 - 1 mg/kg Temgesic 72 hours after the surgery.

### Immunofluorescence staining

Animals were anesthetized under isoflurane, then perfused with 50 ml of cold PBS and 4% paraformaldehyde (PFA). The brains were kept in 4% PFA for 24 hours and then dehydrated in 30% sucrose until they sank. The brains were embedded in OCT and frozen in cold isopentane (2-methylbutyrate) for 2 min. The samples were stored in -80 °C until further use. Frozen brains were cut in a cryostat into 30 μm of free-floating sections and maintained in cryopreservation medium (10 x PBS, ethylene glycol and glycerol, at a ratio of 8:1:1). The sections were washed 4 x 5 min in 1 x PBS, then incubated with 5% normal goat or donkey serum in 0.25% Triton-X100 PBS solution for 1 hour to block the unspecific signal. After the blocking, the sections were incubated overnight at 4°C with unconjugated primary antibodies (listed in **Suppl. Table 3**) in 2.5% normal goat or donkey serum 0.25% Triton-X100 PBS. The next day, the sections were washed 3 x 5 min in 1 x PBS. Secondary antibodies at a concentration of 1:500 were applied to the sections in 1% normal goat or donkey serum in 0.25% Triton-X100 PBS for 2 hours at room temperature. After washing 3 x 5 min with PBS, the sections were stained with 300nM DAPI for 3 min, washed again and covered with Dako Immunofluorescence mounting medium (S302380-2, Agilent). The sections were imaged with either Carl Zeiss Confocal microscopy LSM 880, while tile scan and z-stack was applied when necessary, or with a Revolve Fluorescent Microscope. The images were analyzed using Z-projections (maximum intensity or average intensity). Regions of interest were quantified under NeuN, taking the corresponding regions of interest (ROI), after thresholding for the fluorescence intensity. Mean fluorescence intensity (MFI) for EGFP signal was quantified under the NeuN ROIs. For RNAscope analysis, the mean fluorescence intensity of the *Cnr1* signal was projected into the ROIs of the nucleus of the cell (50 cells were quantified per injection side). All the values were corrected for unspecific background by subtracting values in unstained areas.

### Design and production of gRNAs

The gRNAs that were selected for targeting the promoter of the *H2-Ab1* locus were either designed with CRISPOR (http://crispor.tefor.net/) or manually using the reference mouse genome UCSC July 2007 (NCBI37/mm9). The sequence of the gRNA for LacZ was taken from Gemberling et al ^47^. Cloning of the gRNA requires the CACC and AAAC nucleotides to the 5’ of the sense and antisense oligonucleotides as specific overhangs for the Bpil (BbsI) sites. The oligonucleotides were annealed using 12.5 ul (100 uM) sense oligo, 12.5 ul (100um) antisense oligo, 5 ul fast digest buffer (Thermo Fisher Scientific) and H2O up to 50 ul. The following program was used for the annealing: 95 °C for 5 min, gradually cool down (1 °C/min ramp) from 94 °C to 4 °C and storage in 4°C. The annealed oligos were cloned into the *pKLV2-U6gRNA5(BbsI)-PGKpuro2AmCherry-W* (#67977, AddGene) plasmid using a single reaction including: 2 ul 10X fast digest buffer, 2 ul (10 mM ATP), 500 ng plasmid, 1ul fast digest BpiI restriction enzyme (Thermo Fisher Scientific), 1.5 ul T4 DNA ligase (Thermo Fisher Scientific), 1ul (10 uM) annealed oligo and H2O up to 20 ul. Incubation for 2 hours followed at 37 °C and inactivation of the enzymes was performed at 80 °C for 5-10 min. The plasmids were used for the transformation of TOP10 Electrocompetent E.coli bacteria. After transformation, colony selection was performed and cloning of the gRNAs was verified by sequencing. The transformed bacteria were cultured overnight at 37 °C in LB medium (Luria-Bertani buljong, pH 7,5, MIK2181-1000) supplemented with 100ug/ml of Ampicillin for at least 16 hours. The bacteria were pelleted, and plasmid extraction was followed using the HiSpeed Plasmid Midi kit (Cat.No./ID: 12643, Qiagen). All the gRNA sequences are listed in **Suppl. Table 2** and the plasmid sequences are listed in **Suppl. Table 4**.

### Transfection and sorting of HEK293T cells

Two plasmids, CAG-SV40-Lox2-STOP-3XFLAG-SV40NLS-dCas9-NLS-DNMT3A-T2-EGFP-WPRE and CAG-SV40-Lox2-STOP-3XFLAG-SV40NLS-dCas9-NLS-DNMT3A-IRES-EGFP-WPRE were tested in the HEK293T cells for the methylation in the *BACH2* locus. Briefly, confluent HEK293T cells were transfected using the Lipofectamine™ 3000 (L3000008, Invitrogen) with 4ug of the dCas9-DNMT3A-T2A-EGFP or the dCas9-DNMT3A- IRES-EGFP plasmids, 1.5 ug of the pCAG-iCre plasmid, and 2 ug of the plasmid that was carrying the gRNA for the *BACH2* locus (g8-RNA-mCherry). HEK293T cells were also transfected with 0.5 ug of single plasmids EGFP or mCherry. The medium of the transfected cells was changed 18h later, and the cells were sorted 3 days later using a SONY SH800 Cell Sorter. The cells were collected in FACS tubes, centrifuged at 350 g for 5 min and the supernatant was discarded. For the DNA extraction of the sorted HEK293T cells, we used the QIAamp DNA micro kit, according to the manufacturer’s instructions (QIAGEN, Cat.Number: 56304).

### Lentiviral production

Plasmids containing the gRNAs having mCherry as a reporter, and two packaging plasmids, psPAX2 (#12260, AddGene) and pMD2.G (#12259, AddGene), were used for the production of lentiviruses in HEK293T cells. The cells were cultured in Dulbecco’s Modified Eagle’s Medium (DMEM) - low glucose with 10% heat-inactivated fetal bovine serum (FBS), 100 Uml Penicillin and 100 μg/ml streptomycin, supplemented with L-glutamine, sodium pyruvate and 2-mercaptoethanol. Upon confluency, HEK293T cells were transfected with 5 μg of the gRNA plasmid, and 1 μg of the packaging plasmids per 6-well plate using Lipofectamine™ 3000 (L3000008, Invitrogen). Lentiviral particles were harvested from HEK293T cells at day 3 and 4 after transduction and they were stored overnight at 4℃ in Lenti-X™ Concentrator (631231, Takara bio) at a 1:3 ratio of concentrator:supernatant volumes. For the concentration of the viruses, centrifugation was performed at 1500 g for 45 minutes at 4℃. Viral preparations were stored at -80C until use.

### Bone marrow cell culture and stimulation

Bone marrow cells from femurs and tibia of 6-9 weeks old animals were plated on 12-well plates at a density of 2×10^5 cells per well in complete RPMI medium supplemented with 20 ng/ml of GM-CSF, using a modified protocol from Helft et al. ^48^. At day 3 of culture, 1ml of medium was added and half of the medium was changed at day 6 and 8 with medium containing 20 ng/ml GM-CSF. The cells were stimulated at day 9 of culture for 6 or 24 hours with 10 ng/ml or 100 ng/ml of LPS (TLR2/TLR4 Agonist - LPS from E. coli O111:B4, InvivoGen) respectively, or 20 ng/ml of murine IFNγ for 24 hours (Perpotech, AF-315-05), a combination of the IFNγ/LPS or left untreated. After the stimulation, the loosely adherent and floating cells were harvested, centrifuged at 350 g for 5 min RT and the cell pellet was lysed with RLT plus buffer supplemented with 40 mM DTT. The adherent cells were washed with 1X PBS, and then lysed immediately into the well with RLT plus buffer supplemented with 40 mM DTT. The samples were stored at -80°C until further use.

### Lentiviral transduction of bone marrow cells

The initial trials of lentiviral transduction were performed at day 2 or 5 of bone marrow cell culture, in which the cells were transduced with lentiviruses carrying the gRNAs for the *H2-Ab1* or a control gRNA for LacZ (**Suppl. Table 4**). The transduction was performed with spinoculation for 1.5 hours at 3000 g at RT. At day 4 (Day-2 transduction) or 7 (Day-5 transduction), complete medium was changed, and at day 6 (Day-2 transduction) and 9 (Day-5 transduction), complete medium was changed with the medium supplemented with 20 ng/ml GM-CSF. For the sorting (at day 9 or 12), the dendritic cell fraction was removed and centrifuged at 350 g for 5 min, while the macrophage layer was incubated with 1X PBS and 5 mM EDTA at 37°C until they detached and proceeded to sorting, where the cells were sorted using a SONY SH800 Cell Sorter. After sorting the cells were lysed in RLT Plus Buffer and were stored at -80℃ until further use.

### Tissue collection, nucleic acid extraction, and gene expression analysis

Animals were perfused with 50 ml of cold PBS and 30 mg of tissue was collected from different organs. The samples were snap frozen on dry ice and kept at -80℃ until further use. Simultaneous DNA and RNA extraction was performed using the AllPrep DNA/RNA Mini Kit from approximately 30 mg of tissue. A TissueLyser LT (QIAGEN) was used for the disruption and homogenization of the samples with the DNA lysed in RLT Plus buffer supplemented with 40 mM DTT according to the manufacturer’s instructions. For the bulk intracerebral injections, up to 5 mg of tissue was isolated from fresh brain tissue after animal perfusion with 50 ml of cold PBS. The dorsolateral part of the striatum was dissected under a stereoscope, and the tissue was frozen on dry ice until further use. DNA/ RNA extraction was followed using the AllPrep DNA/RNA Micro kit (Qiagen, 80284). RNA concentration was measured with the QIAexpert Instrument (QIAGEN). A range of 50-200 ng of RNA was used for cDNA synthesis with the iScript cDNA Synthesis Kit (BIO-RAD). RT- qPCR was performed with the iQ SYBR Green Supermix kit (BIO-RAD) and the quantification with Biorad CFX Maestro. Samples with Ct values differing more than 1 Ct cycle between experimental duplicates were removed from the analysis. The relative expression (2^-ΔΔCt) was determined by Ct values of the *dCas9-DNMT3A* gene relative to the *Gapdh* or *Hprt* housekeeping genes.

### Pyrosequencing

Genomic DNA was treated with sodium-bisulfite that converts the unmethylated cytosines to uracils, while it preserves the methylated cytosines, during amplification using primers designed to anneal to bisulfite-converted DNA sequence. In total 200 ng of genomic DNA from different tissues was converted using the EZ DNA Methylation-Gold Kit (Zymo Research Europe, Freiburg, Germany) according to the manufacturer’s protocol. Bisulfite-converted DNA (10 ng) was amplified by PyroMark PCR kit (Qiagen, 978703, Hilden, Germany) according to the manufacturer’s instructions.

The PCR was performed using 5 pmol of forward and 5′-biotinylated reverse primers with the following cycles: initial denaturation for 15 min at 95^◦^C; 50 cycles of 30 s at 94°C, 30 s at 48°C (*BACH2*) and 30 s at 72°C; final extension for 10 min at 72^◦^C. For all the other genes, *H2-Ab1*, *Il6* and *Cnr1*, PCR conditions were the following: 15 min at 95°C; 50 cycles of 30 s at 94°C, 30 s at 55°C, and 30 s at 72°C; final extension for 10 min at 72°C. Single band amplification was checked on 1% agarose gel and PCR products were sequenced by Pyromark Q96 pyrosequencing system (Qiagen) using PyroMark Gold 96 reagent kit (Qiagen, 972804) or a PyroMarkQ48Autoprep pyrosequencing using the PyroMark Q48 Advanced CpG Reagents (Qiagen, 974022). All pyrosequencing PCR primers and probes were designed by PyroMark Design software (Qiagen) and are listed in the **Suppl. Table 2**.

### Fluorescent In situ hybridization

The animals were sacrificed with cervical decapitation, brains were removed and snap frozen in cold 2-methylbutane on dry ice. Sections with 12 um thickness were mounted onto Superfrost slides, kept at -80°C until further use. At the day of RNAscope, the frozen tissue was immersed into 10% Neutral Buffered Formalin (NBF) for 30 min, the tissue was then dehydrated with increasing concentration of ethanol, and treated with Protease-free solution (PretreatProTM) for 30 min at RT. Afterwards, the tissue was hybridized with the probes (RNAscope™ Probe-Mm-Cnr1-O1-C3, Cat No. 457341-C3 (TSA Vivid Fluorophore 650), Mm-PPIB-C2 #313911 (TSA Vivid Fluorophore 570), positive control, or DapB-C1 #310043 negative control (TSA Vivid Fluorophore 650), for 2 hours at 40°C, followed by amplification with AMP-1 and AMP-2 for 30 min, and AMP-3 for 15 min at 40°C. The tissue was incubated with HRP-C1, HRP-C2, or HRP-C3 for 15 min at 40°C, and TSA (Tyramide Signal Amplification) was applied for 30 min at 40°C. After the development of the probe signal, the sections were incubated with anti-TdTomato (1:200, A121690) overnight in 5% Normal Donkey Serum in 0.25% T-TBS. After the incubation, the sections were washed 3 X 5 minutes in PBS, and donkey anti-goat Alexa Fluor 555 antibody was applied for 2 hours (Catalog # A-21432, 1:500). Finally, the sections were counterstained with DAPI, mounted with aqueous mounting medium (Vectashield, H-1900-2) and covered with coverslips. Fluorescence images were acquired with a Carl Zeiss LSM 880 confocal microscope, using tile scan and z-stack acquisition with step size of 1 μm ^49^.

### Genome-wide DNA methylation profiling

In total 500 ng of genomic DNA from bulk brain and spleen tissue was diluted in 45 ul of low EB buffer 10mM Tris-Cl, pH 8.5 and sent for the Infinium Mouse Methylation BeadChip array profiling using a 12 sample array, performed in BEA (Bioinformatics and Expression Analysis Core Facility). Brain and spleen samples were run separately in two arrays. Data were analyzed using R version (4.5.0) and the SeSAMe package (version 1.28.0). IDAT files were imported to R, masking of the probes was performed by inferring the strain (C57BL/6J), and later preprocessing was performed using the functions: pOOBAH, noob and dyeBiasNL ^50^. Normalized beta values were extracted and comparison performed using linear modelling in R (Δβ < 0.05 and Δβ > 0.05, and p<0.001). Annotation of the probes was done using Illumina’s manifest MM285.mm10.manifest (https://github.com/OluwayioseOA/Aclust2.0/blob/main/MM285.mm10.manifest.csv). Normalized beta values were converted to z-scores using the https://heatmapper.ca.

### Flow cytometry

All antibodies used for flow cytometry are listed in **Supplementary Table 3**. The assessment of the myeloid and lymphoid (T, B and NK cells) compartment was conducted in single cell suspensions obtained from lymph nodes and spleens. The spleens were smashed, followed by red blood cells removal using ACK lysis buffer (Gibco, A1049201) for 5 min in room temperature. Cells were preincubated with Fc block (1:1000) and Yellow-Dead Dye (1:500, Invitrogen, L34959) or NearIR-Dead Dye (1:500, L34976) for 20 min at 4°C in PBS, followed by the incubation with surface marker for the different panels. CountBright Absolute Counting Beads (Thermo Fisher Scientiifc, C36950) were added to the samples for the absolute quantification of the cell numbers in the corresponding organ. The samples were acquired with LSR Fortessa (BD). Data analysis was performed with FlowJo v10.10 (BD Life Sciences).

### Experimental autoimmune encephalomyelitis

Mice were immunized subcutaneously at the tail base with a 100ul emulsion composed of 20-30ug of recombinant myelin oligodendrocyte glycoprotein (MOG), and Complete Freud’s Adjuvant that contained 100 ug inactivated MTB (Chondrex Inc) prepared using the POWER-Kit™ (BTB Emulsions). At day 0 and 2 after immunization, Pertusssis toxin was injected intraperitoneally to the animals (200ng to 200ul, Millipore Corp). The development of the disease was assessed from day 7 until the termination of the experiment and the following scoring scale was used: 0, healthy; 1, tail weakness or tail paralysis; 2, hind leg paresis or hemiparesis; 3, hind leg paralysis or hemiparalysis; 4, tetraplegy; and 5, death. Cumulative score and Maximum EAE score were also assessed in the animals.

### Statistical analysis

Statistical analysis was performed with Prism 10 (Version 10.2.3) (GraphPad, San Diego, USA). The Shapiro Wilk Normality test for small sample numbers was performed before selecting to run parametric or non-parametric tests for group comparisons. When more than two groups were compared for variances that did not follow the Gaussian distribution, non-parametric Kruskal Wallis test with Dunn’s correction for multiple comparisons was performed. Unpaired and paired t-test, as appropriate, was used to compare normally distributed variables between two groups.

## Supporting information

Suppl. Figures_Kalomoiri et al

## Data availability

The bioinformatic datasets for the Infinium Mouse Methylation BeadChip arrays are available in the NCBI Gene Expression Omnibus (GEO) under accession codes GSE (GSE325144, GSE325146).

## Code availability

Custom code developed for this study, along with a valid license, is available via GitLab at https://gitlab.com/jagodiclab/mouse-methylation-epic.

## Acknowledgements

We wish to acknowledge Dr. Alexander Espinosa and Dr. William A. Nyberg for their valuable input, and Dr. Andre Ortlieb Guerreiro-Cacais and Annika van Vollenhoven for their assistance with flow cytometry analysis and sorting. We thank Dr. Sho Oasa for his assistance with confocal microscopy. We acknowledge the National Academic Infrastructure for Supercomputing in Sweden (NAISS) at UPPMAX, funded by the Swedish Research Council through grant agreement No. 2022-06725 for providing computational resources, the National Genomics Infrastructure (NGI) in Stockholm funded by Science for Life Laboratory, the Knut and Alice Wallenberg Foundation and the Swedish Research Council, and the Center for Molecular Medicine (CMM) for providing research facilities, including the KI Gene and Flow Cytometry Core, among others.

This work was supported by grants from the European Union Horizon 2020 Research and Innovation Programme/European Research Council Consolidator (grant Epi4MS, No. 818170), Swedish Research Council, Swedish Association for Persons with Neurological Disabilities, Swedish Brain Foundation, Swedish MS Foundation, Karolinska Institutet, StratNeuro, and the Knut and Alice Wallenberg Foundation. M. Pahlevan Kakhki was supported by the McDonald fellowship from the Multiple Sclerosis International Federation (MSIF) and the Innovative Medicines Initiative 2 Joint Undertaking (JU) under grant agreement No. 875510 (**Suppl. Table 5**). L. Kular was supported by fellowship from the Margaretha af Ugglas Foundation.

## Author contributions

MJ conceived the study. MK, LK, MPK and MJ designed the experiments. MK performed and analyzed all experiments with help from CS, SV, AC, AT, AK, CRP, MN and MPK. MK, LK, MPK and MJ wrote the manuscript. LK, MPK, PS and MJ supervised the project. All authors read and approved of the final manuscript.

## Additional information

Supplementary information

