## Supplementary material for "Transgenic dCas9-DNMT3A mouse lines enable *ex vivo* and *in vivo* methylation editing with locus-dependent transcriptional effects": Suppl. Figures_Kalomoiri et al

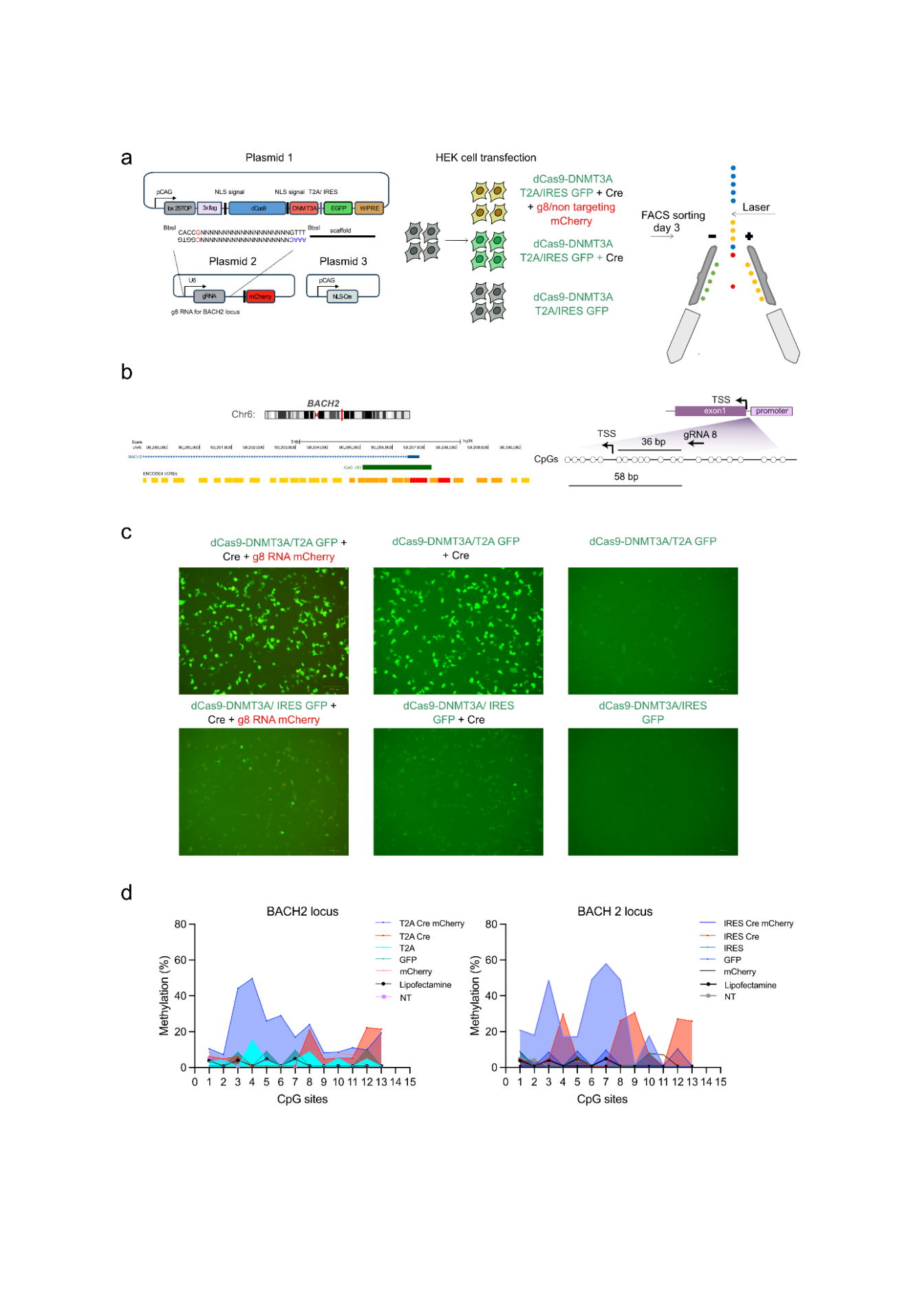
(Kalomoiri et al.)

**Supplementary Fig. 1** I **Investigation of the impact of two different dCas9-DNMT3A constructs on the methylation profile in the *BACH2* promoter.** **a.** Graphical presentation of the three different plasmids used for the transfection of HEK293T cells, encoding the dCas9-DNMT3A epimodifier with GFP reporter, the *BACH2 or NTC*-targeting gRNA with mCherry reporter, and the Cre recombinase. HEK293T cells were transiently transfected with combination of plasmids and sorted 3 days post-transfection. **b**. Representation of the *BACH2* locus (chr6:89,926,528-90,296,843 370,316) from the UCSC browser, utilizing relevant tracks and the relative positions of the CpGs from the Transcritpional Start Site (TSS). **c.** Representative images of transfected cells 24 hours after transfection. Methylation analysis of the *BACH2* promoter, covering 13 CpGs downstream of the gRNA binding site, following transfection with **d.** dCas9-DNMT3A-T2A-GFP and dCas9-DNMT3A-IRES-GFP.

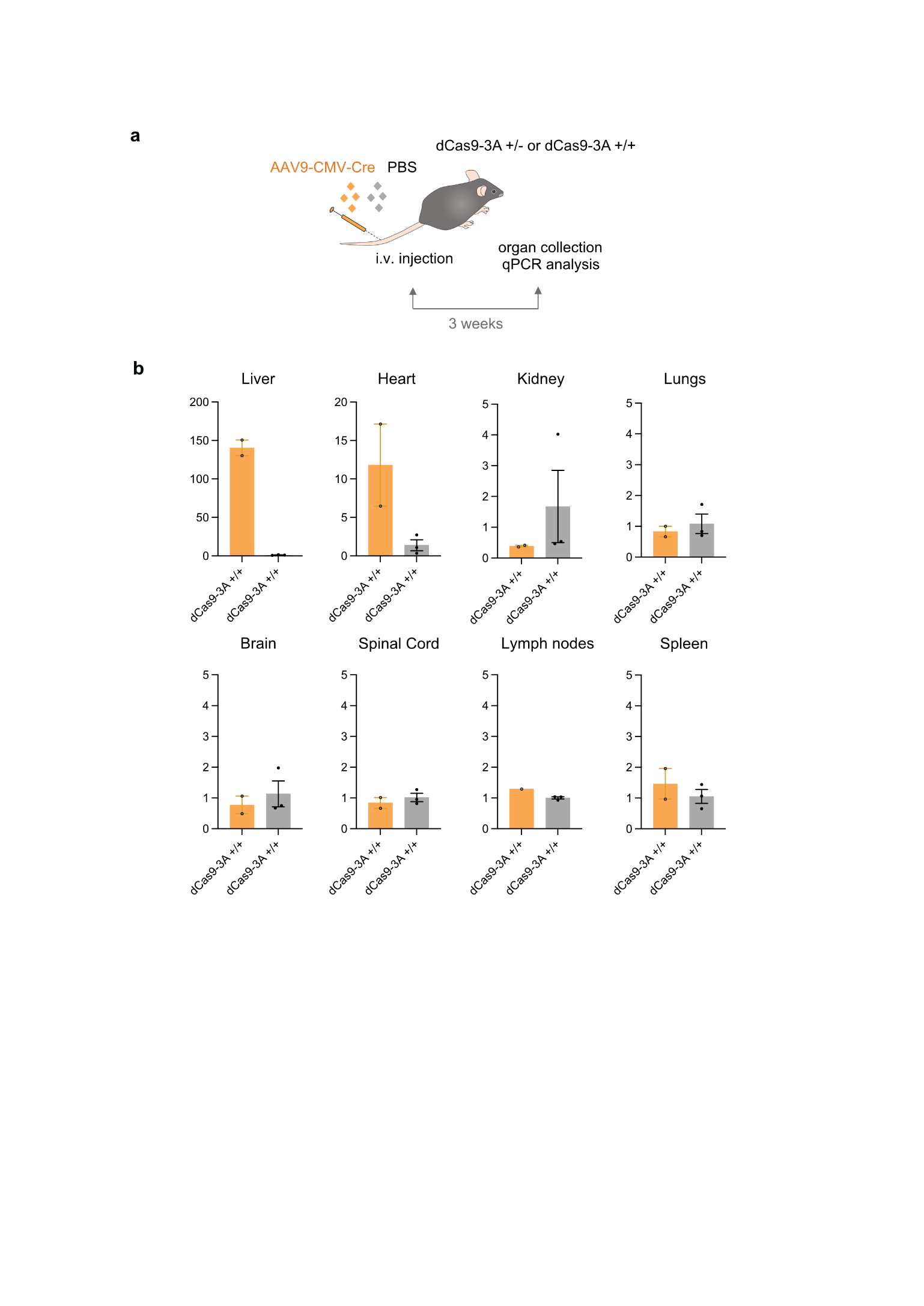

**Supplementary Fig. 2** I **Induction of the cassette in several organs of the dCas9-3A +/+ though intravenous injection of AAV9-CMV-Cre viruses.** **a.** Schematics of the experimental design of intravenous injections with AAV9-CMV-Cre viruses or PBS control. **b.** Fold change of *dCas9-DNMT3A* expression (normalized to Hprt) assessed by qPCR in the organs harvested from heterozygous dCas9-DNMT3A animals injected with AAV9 or PBS. Data are represented as mean ± S.E.M. with n=1-3 per group.

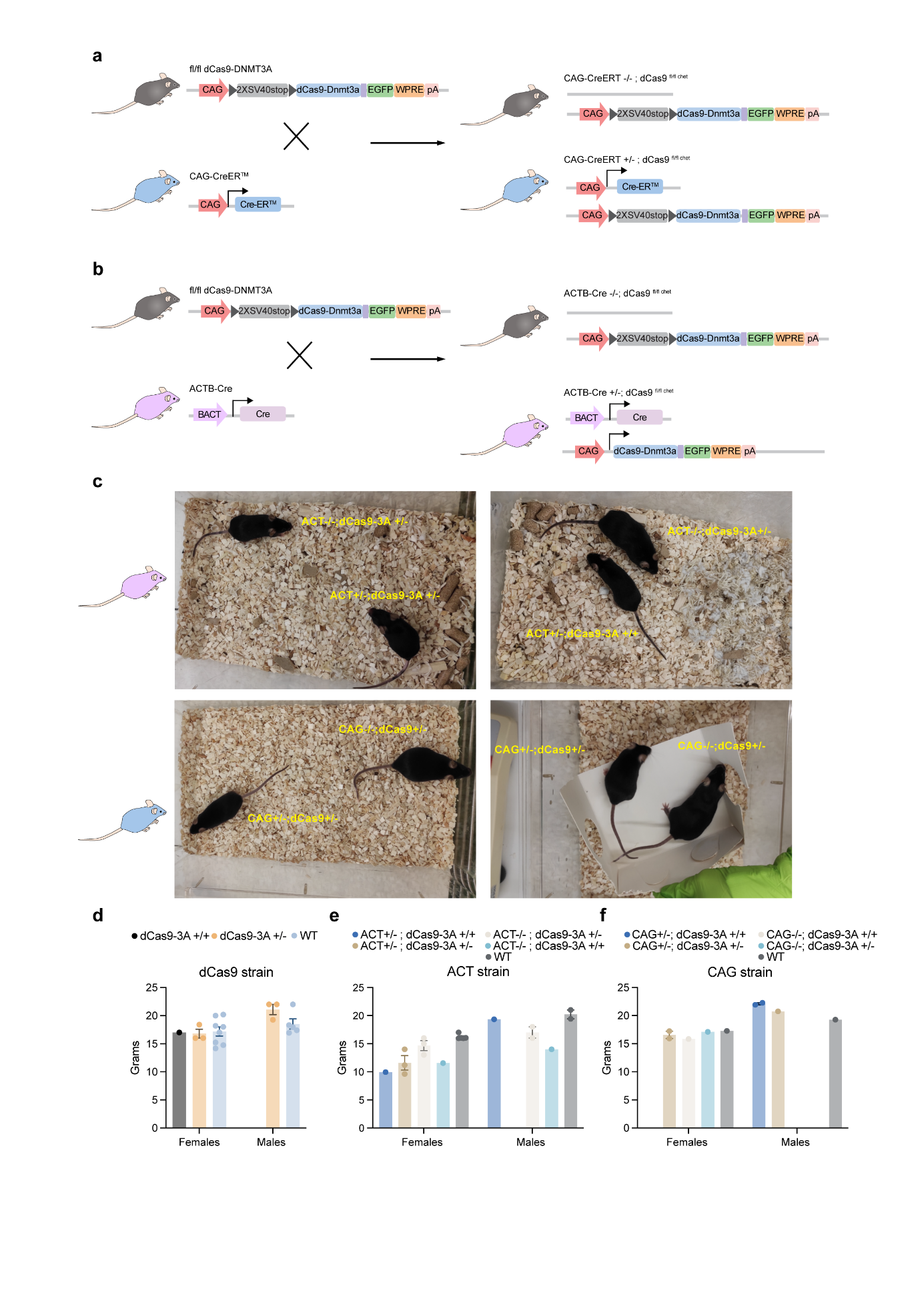

**Supplementary Fig. 3** I **Generation of the conditional and constitutive expressing lines.** Generation of **a.** the conditional CAG-CreERT+/-;dCas9fl/fl mouse line and **b.** the constitutive ACTB-Cre+/-; dCas9fl/fl line by crossing the original dCas9-DNMT3Afl/fl line with CAG-CreERT+/- and ACTB-Cre+/- mice, respectively. The floxed animals from respective breedings were used as littermate controls. **c.** Representative images of the body size of adult ACT+-; dCas9-3A+/- and CAG+/-; dCas9-3A+/- animals in comparison to littermate controls. **d-f**. Weight of at 5-6 weeks of age of **d.** the original dCas9, **e.** ACT;dCas9-3A, and **f.** CAG;dCas9-3A strain animals.

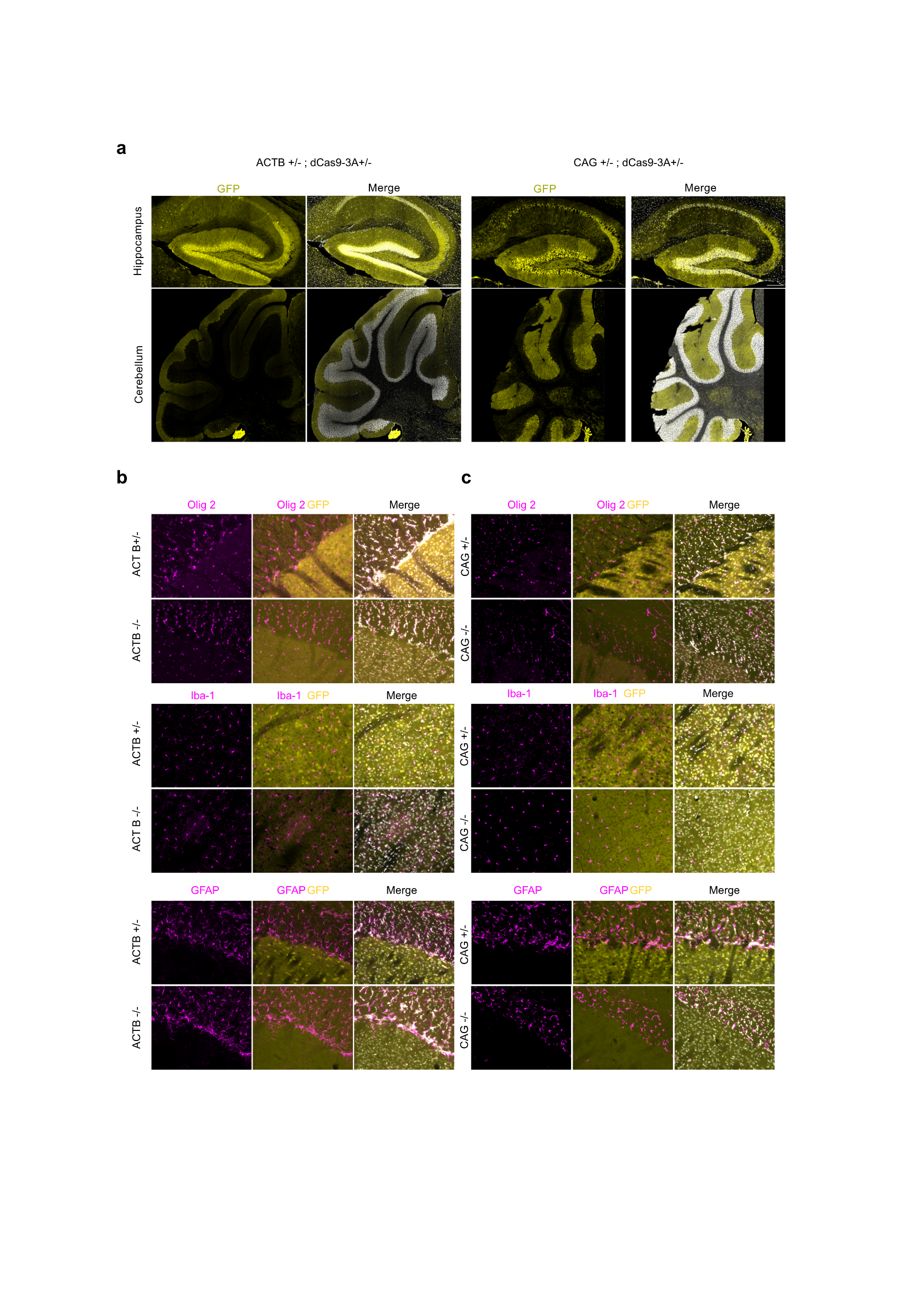

**Supplementary Fig. 4** I **Assessment of the EGFP expression in different CNS population in ACTB;dCas9 and CAG;dCas9 animals. a.** Confocal images of hippocampus and cerebellum of ACTB+/-;dCas9-3A +/- and CAG+/-;dCas9-3A +/- animals. Fluorescence microscope images of EGFP localization in oligodendrocytes (Olig2), microglia (Iba-1) and astrocytes (GFAP) in corpus callosum and striatum of **b.** ACT-B+/-;dCas9-3A +/- and **c.** CAG+/-;dCas9-3A +/- animals, and respective floxed controls. Scale bar: 200um (hippocampus) and 200um (cerebellum) in a,100um in b.

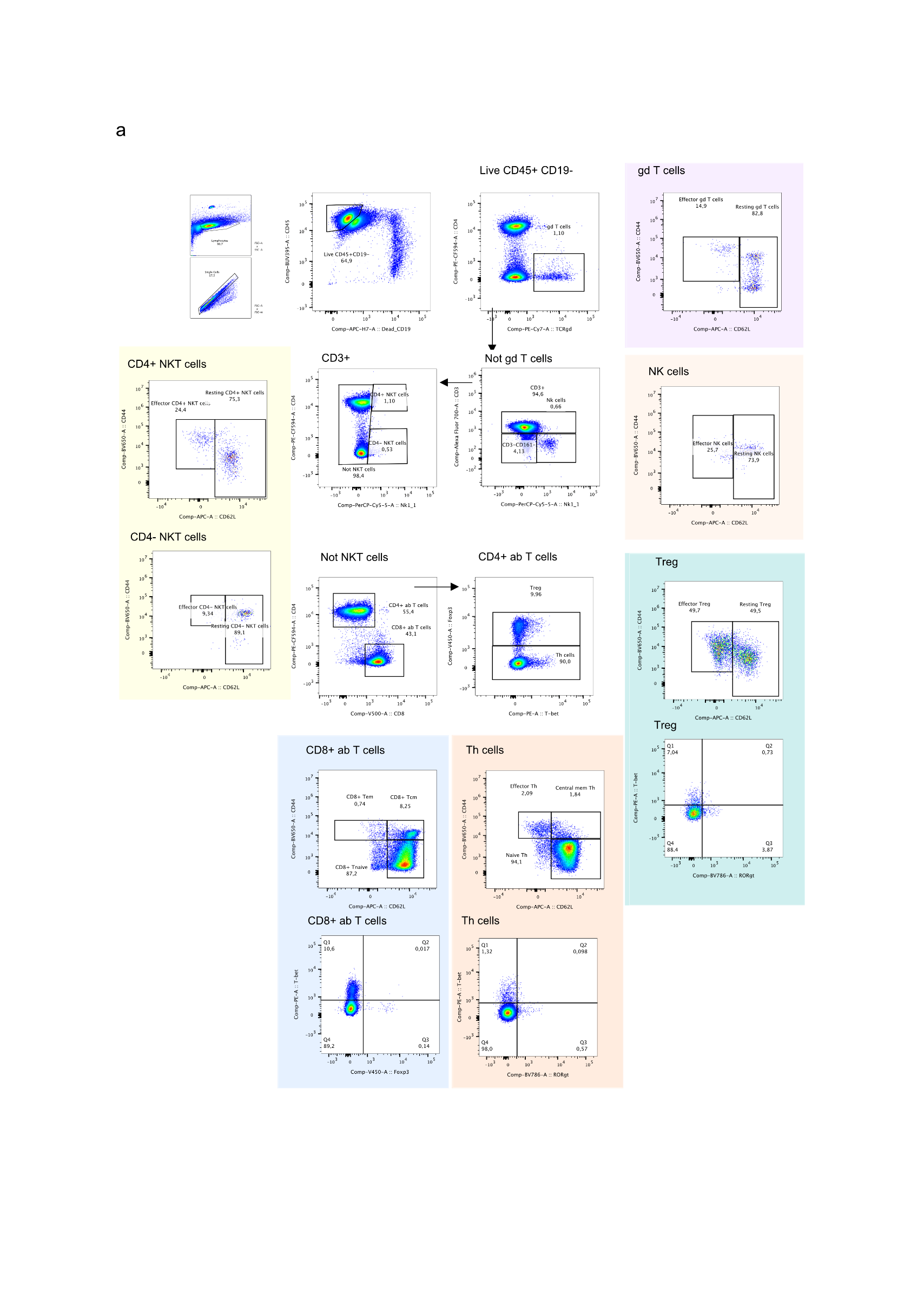

**Supplementary Fig. 5** I **Gating strategy for T and NK cell compartments**. Representative example of flow cytometry gating strategy for T and NK cell populations from the lymph nodes of naive animals.

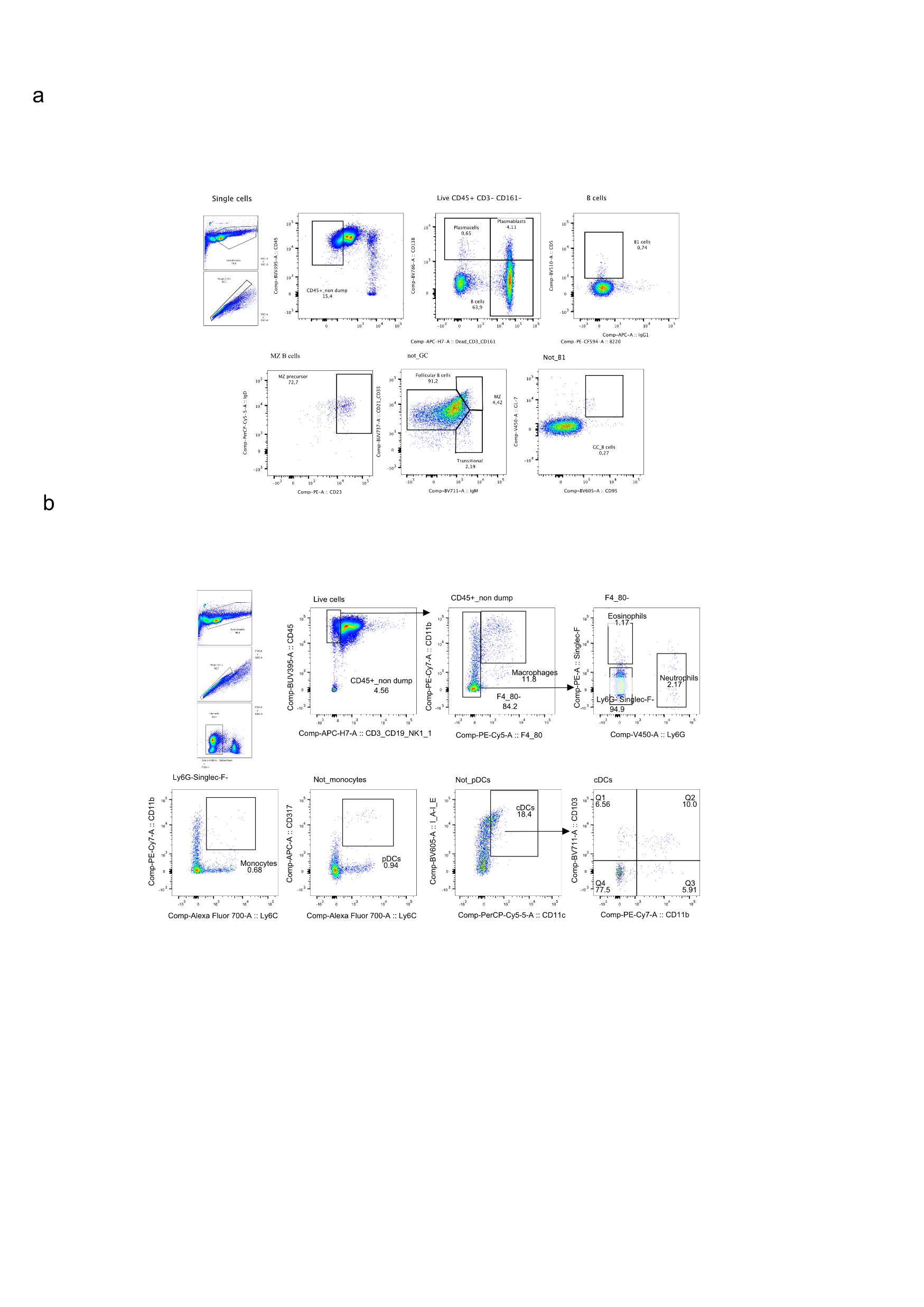

**Supplementary Fig. 6** I **Gating strategy for B and myeloid cell compartments**. Representative example of flow cytometry gating strategy for **a.** B cell, and **b.** myeloid cell populations from the lymph nodes of naïve animals.

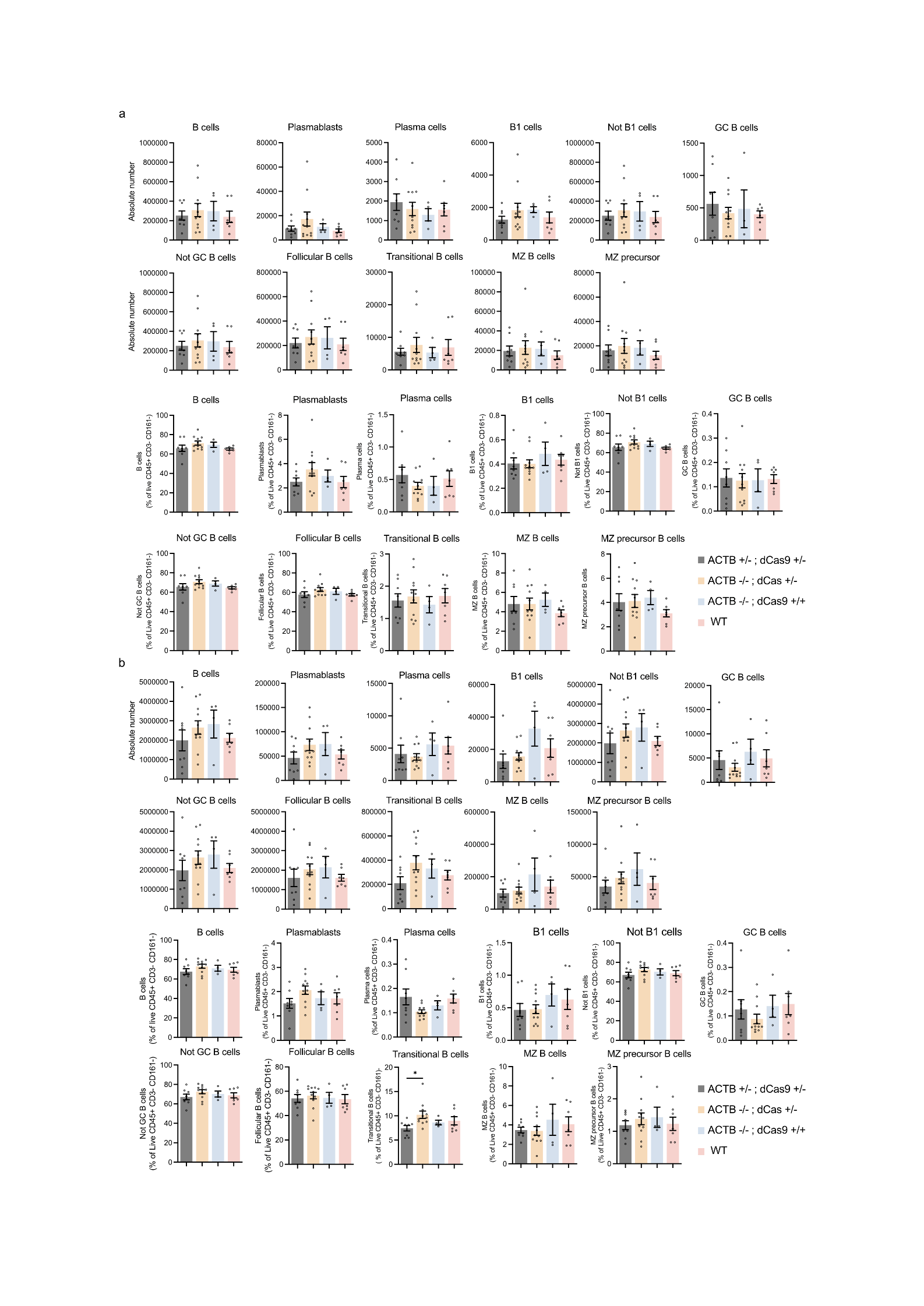

**Supplementary Fig. 7** I **The B cell compartment.** The absolute numbers and percentages of B cell subsets in **a.** lymph nodes and **b.** spleen of naive animals (n=8 for ACTB+/-; dCas9-3A +/-, n=11 for ACTB-/-; dCas9-3A +/-, n=4 for dCas9-3A +/+, n=7 for WT animals). Data are represented as Mean ± S.E.M. One-way ANOVA Kruskal Wallis test with Dunn’s multiple corrections tests was performed. *, p<0.05, non-significant p-values are not represented.

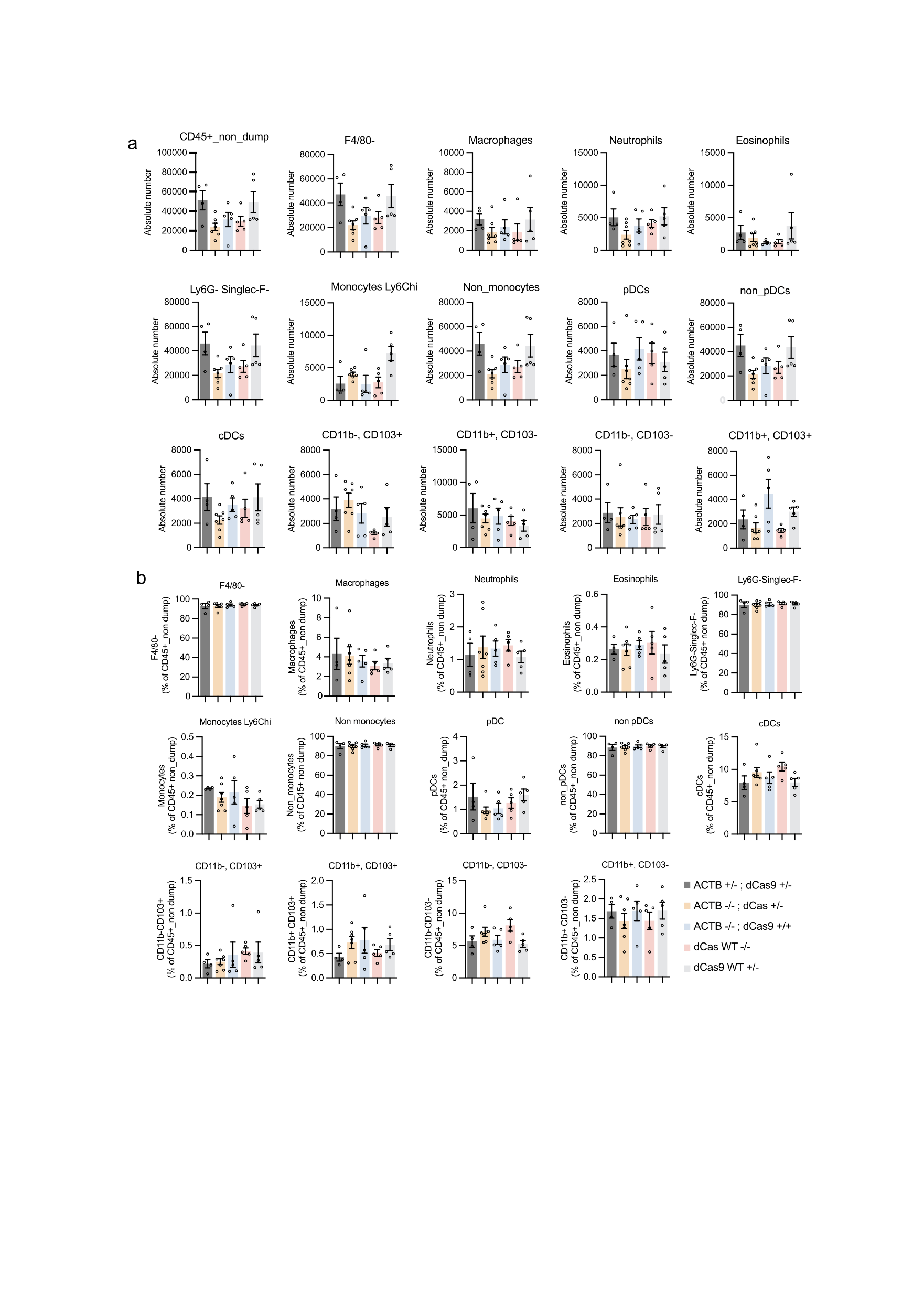

**Supplementary Fig. 8** I **The lymph node myeloid compartment.** The **a.** absolute numbers and **b.** percentages of myeloid cell subsets in lymph nodes of naive animals (n=4 for ACTB+/-; dCas9-3A +/-, n=7 for ACTB-/-; dCas9-3A +/-, n=5 for dCas9-3A +/+, n=5 for WT animals). Data are represented as Mean ± S.E.M. One-way ANOVA Kruskal Wallis test with Dunn’s multiple corrections tests was performed. Non-significant p-values are not represented.

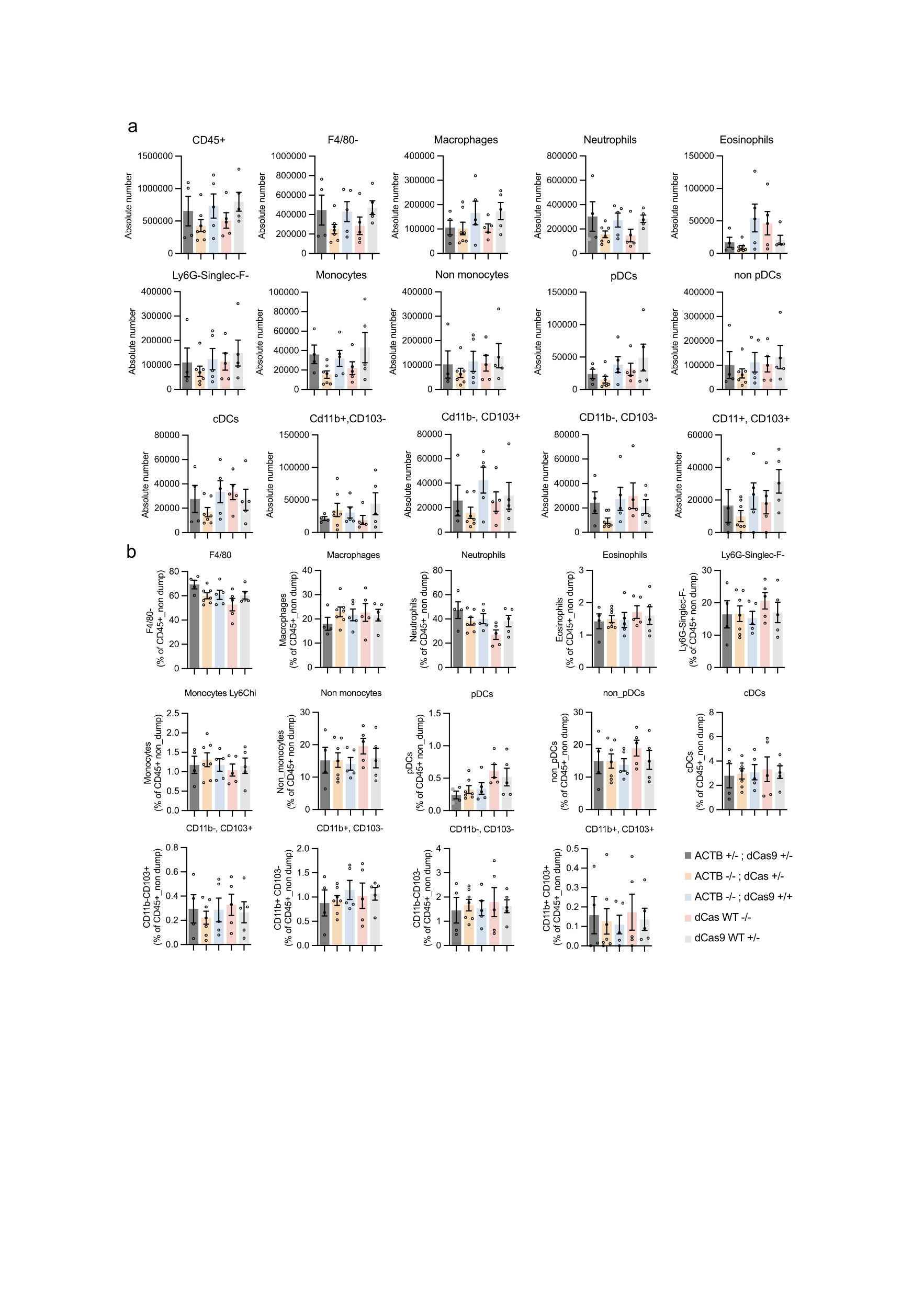

**Supplementary Fig. 9** I **The spleen myeloid compartment.** The **a.** absolute numbers and **b.** percentages of myeloid cell subsets in spleen of naive animals (n=4 for ACTB+/-; dCas9-3A +/-, n=7 for ACTB-/-; dCas9-3A +/-, n=5 for dCas9-3A +/+, n=5 for each WT animal category). Data are represented as Mean ± S.E.M. One-way ANOVA Kruskal Wallis test with Dunn’s multiple corrections tests was performed. Non-significant p-values are not represented.

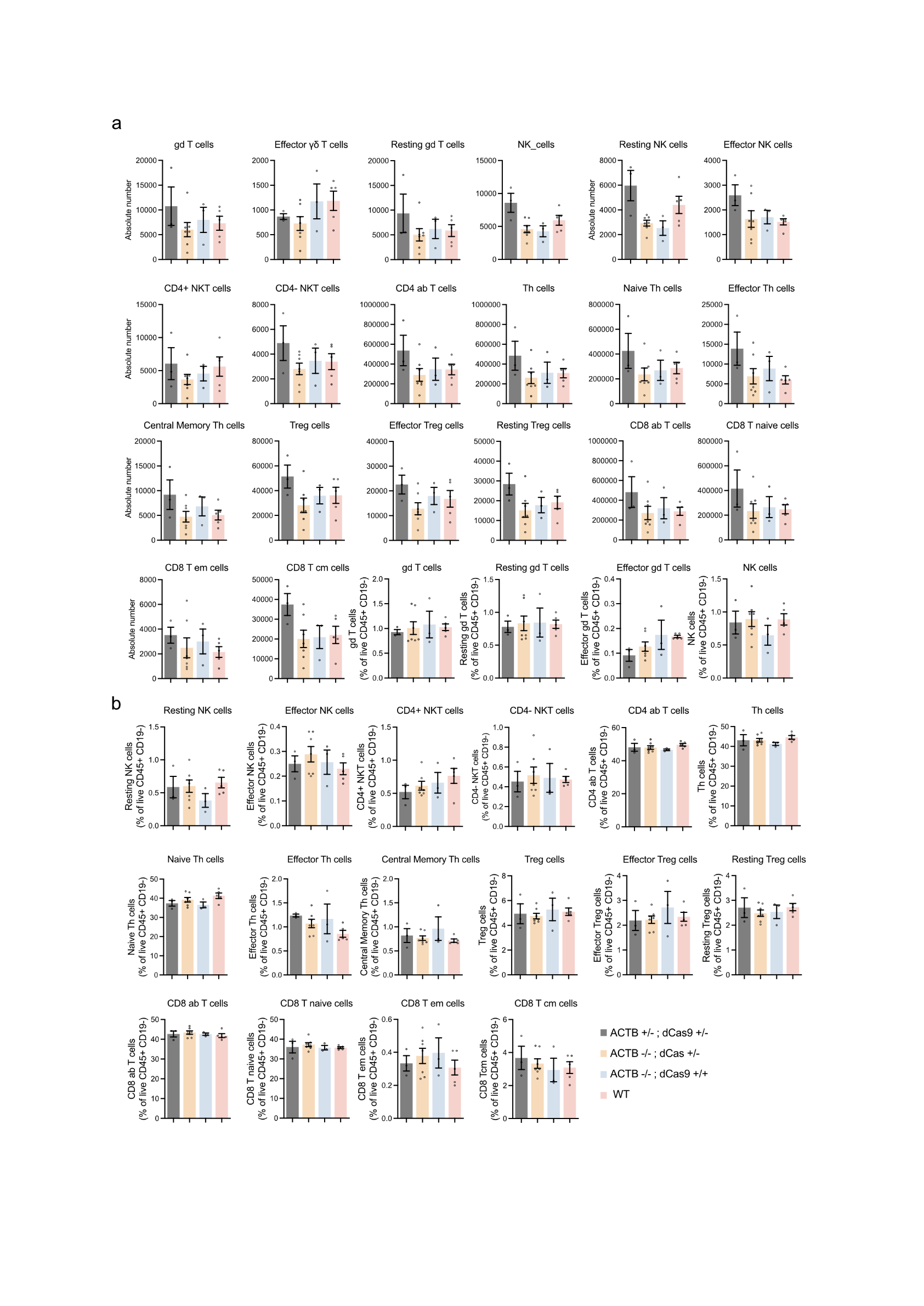

**Supplementary Fig. 10** I **The lymph node T and NK cell compartment**. The **a.** absolute numbers and **b.** percentages of T and NK cell subsets in lymph nodes of naive animals (n=3 for ACTB+/-; dCas9-3A +/-, n=7 for ACT-/-; dCas9-3A +/-, n=3 for dCas9-3A +/+, n=5 for WT animals). Data are represented as Mean ± S.E.M. One-way ANOVA Kruskal Wallis test with Dunn’s multiple corrections tests was performed. Non-significant p-values are not represented.

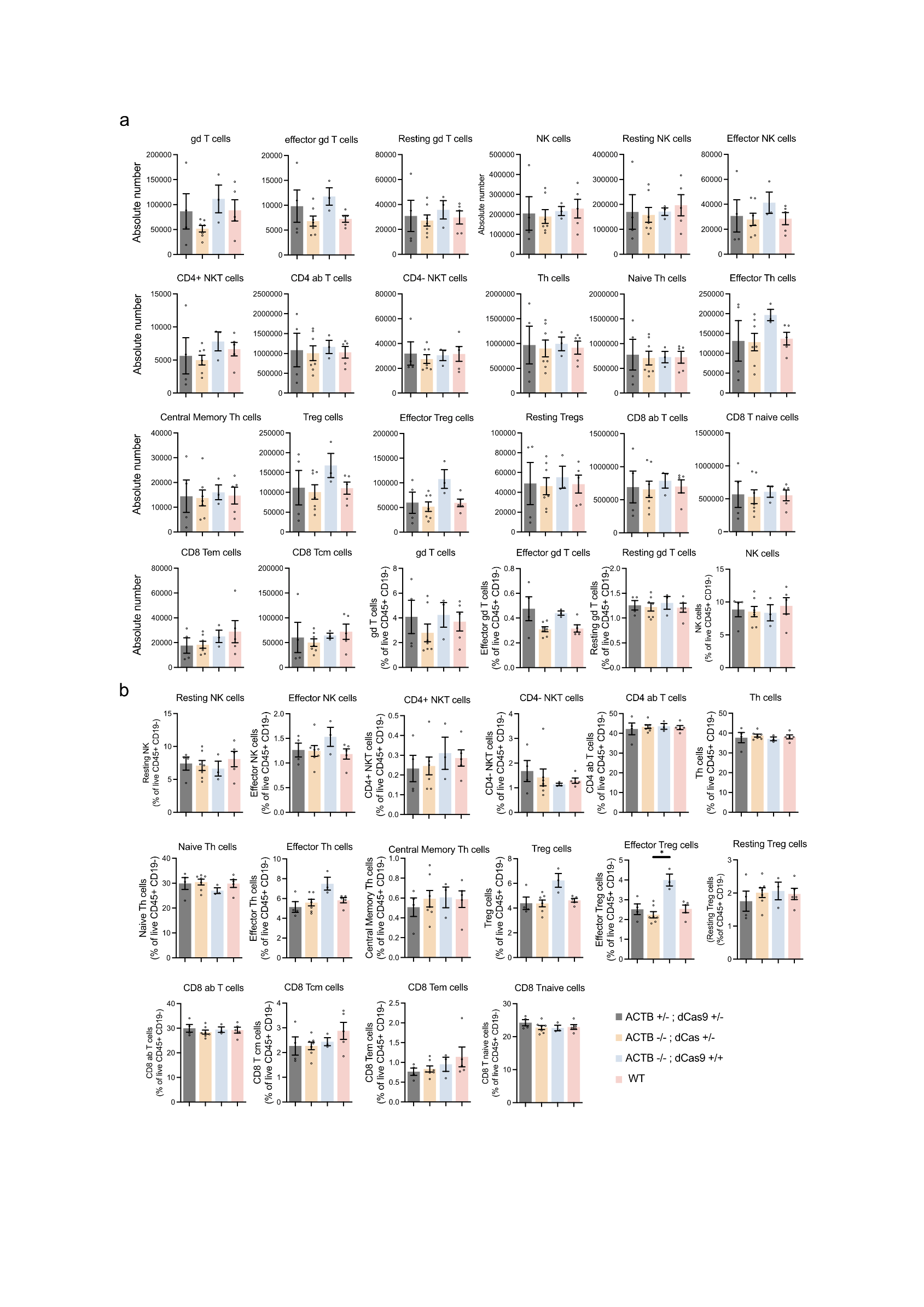
**Supplementary Fig. 11** I **The spleen T and NK cell compartment**. The **a.** absolute numbers and **b.** percentages of T and NK cell subsets in spleen of naive animals (n=4 for ACT+/-; dCas9-3A +/-, n=7 for ACT-/-; dCas9-3A +/-, n=3 for dCas9-3A +/+, n=5 for WT animals). Data are represented as Mean ± S.E.M. One-way ANOVA Kruskal Wallis test with Dunn’s multiple corrections tests was performed. Non-significant p-values are not represented.

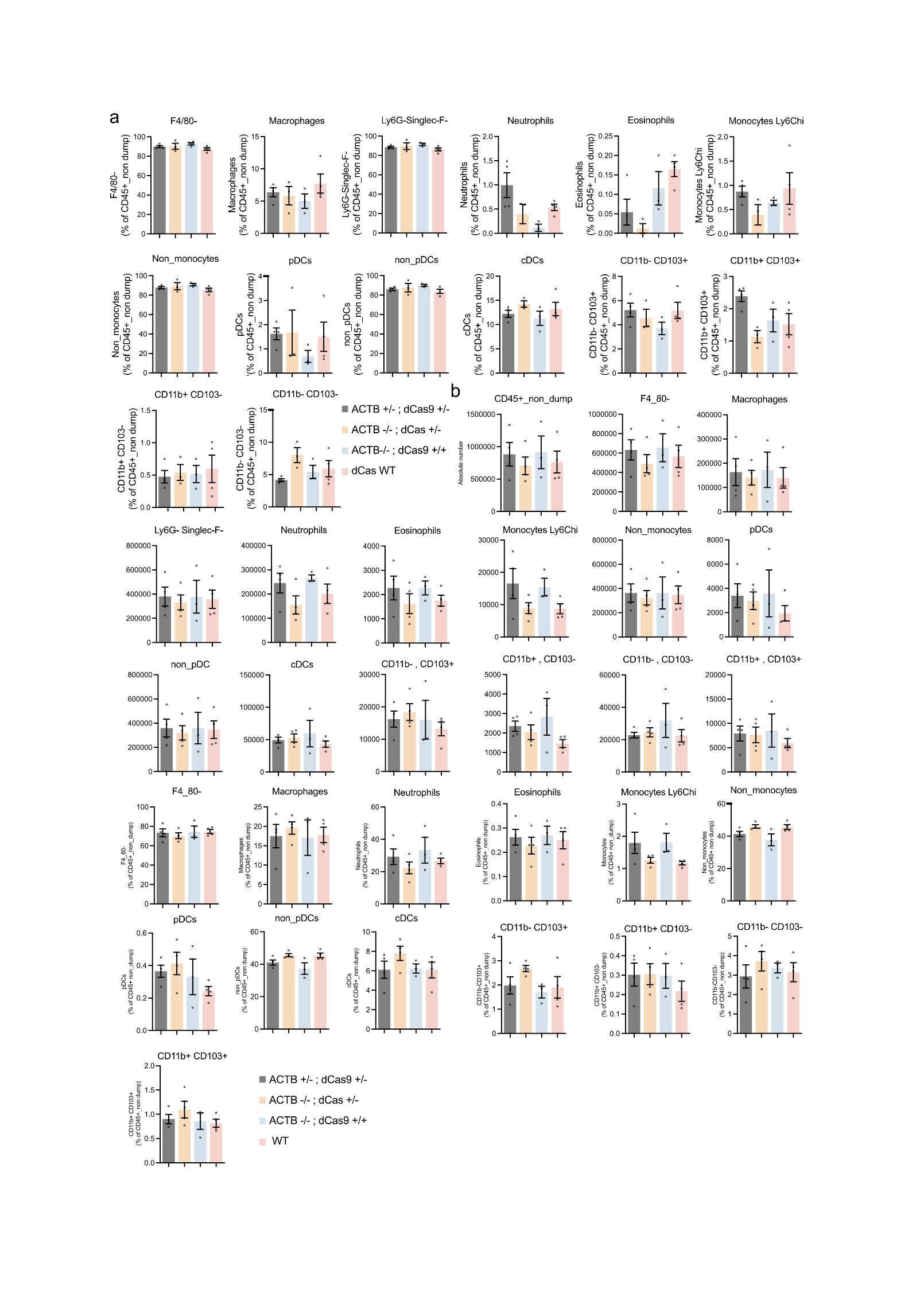
**Supplementary Fig. 12** I **The myeloid compartment in older animals.** The absolute numbers and percentages of myeloid cell subsets in **a.** lymph nodes and **b.** spleen of naive 33-week-old animals (n=4 for ACTB+/-; dCas9-3A +/-, n=3 for ACTB-/-; dCas9-3A +/-, n=3 for dCas9-3A +/+, n=4 for WT animals). Data are represented as Mean ± S.E.M. One-way ANOVA Kruskal Wallis test with Dunn’s multiple corrections tests. Non-significant p-values are not represented.

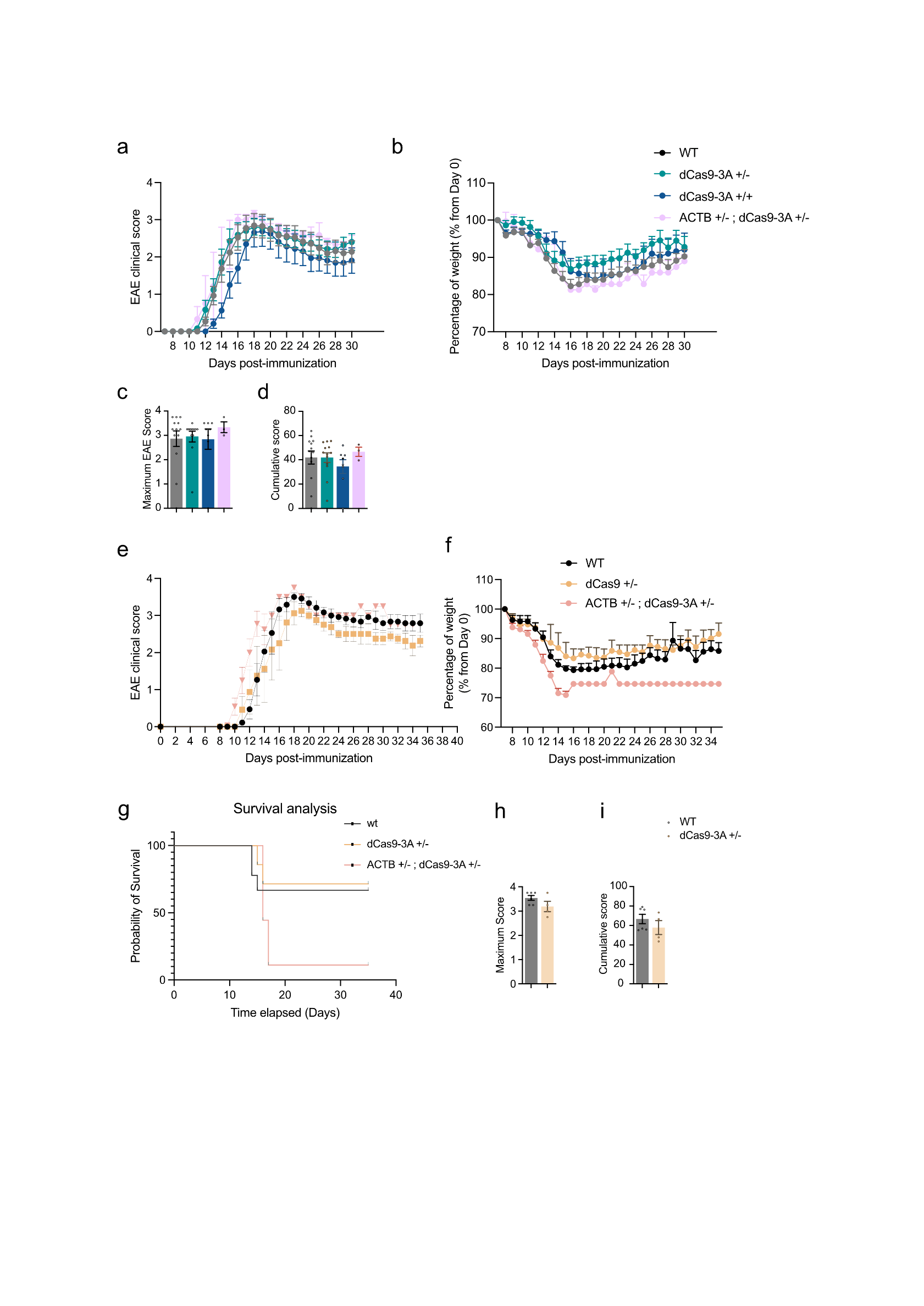

**Supplementary Fig. 13** I **Development and characteristics of Experimental Autoimmune Encephalomyelitis.** Animals were immunized with recombinant myelin oligodendrocyte glycoprotein in adjuvant and clinical signs of Experimental Autoimmune Encephalomyelitis (EAE) disease and weight were followed until day 35 post-immunization in **a.** 11-15 weeks old and **e.** 39-44 weeks old for ACTB+/-;dCas9-3A +/- animals and 25-35 weeks old for the dCas9-3A strain (n=3-13 per group, representative of two experiments in **a,b** and one experiment in **e,f**). Statistical analysis was performed using Kruskal-Wallis test followed by Dunn’s multiple correction test. Data is expressed by mean ± SEM. Non-significant values are not presented.

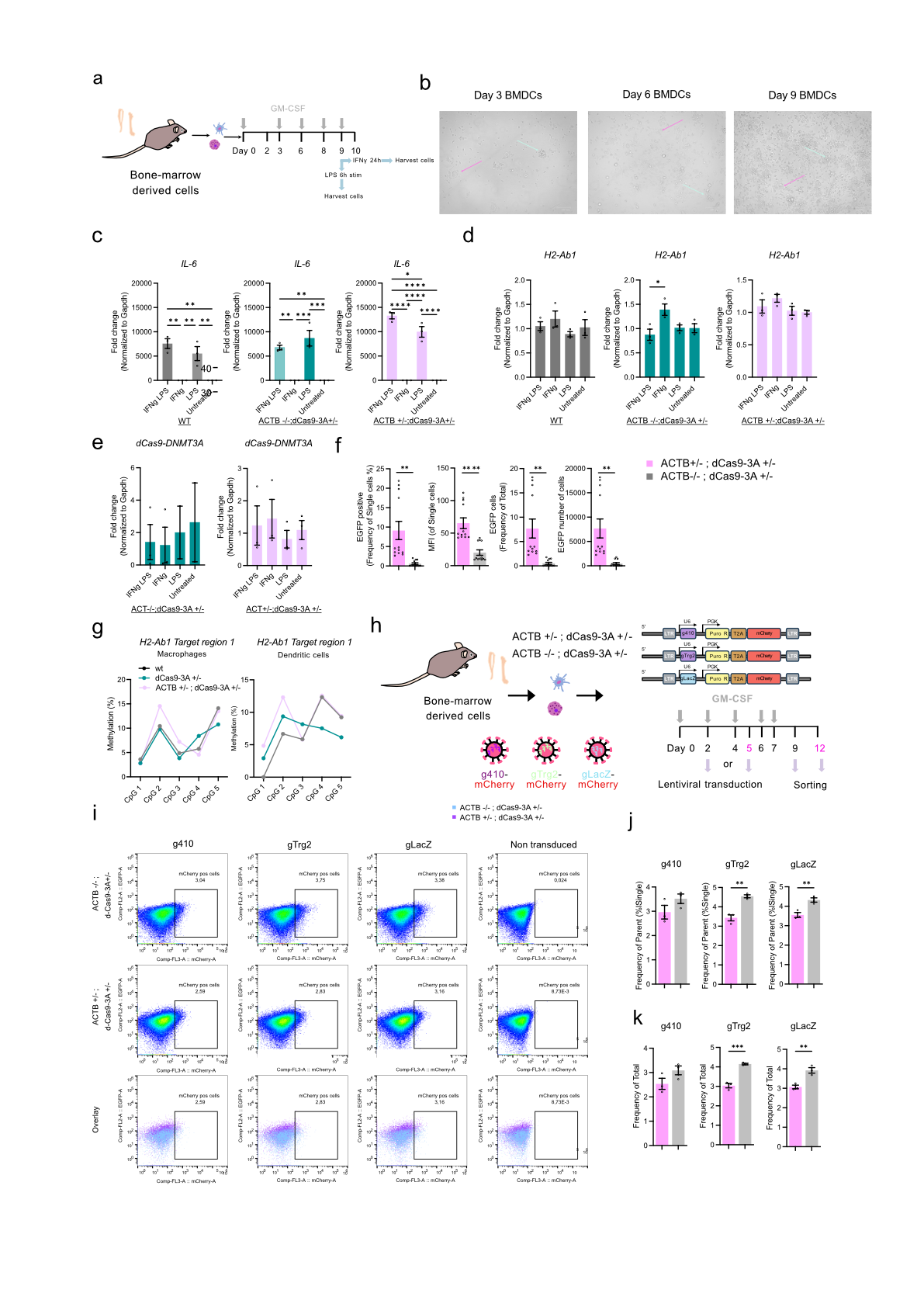

**Supplementary Fig. 14 I Establishment of relevant assays in *ex vivo* bone-marrow derived-dendritic cells and macrophages.** **a.** Experimental procedure for the assessment of DNA methylation upon application of different stimuli. **b.** Microscopy images from the bone-marrow derived dendritic cells and macrophages. **c-e.** *IL-6, H2-Ab1* and *dCas9* expression estimation upon different inflammatory conditions after 6 hours of stimulation in different genetic background animals (n=3 per condition, except the dCas9-3A where n=2 in LPS and untreated). **f.** EGFP expression of lentivirally transduced and sorted cells of ACTB+/-; dCas9-3A +/- and ACTB-/-; dCas9-3A +/- animals. **g.** Assessment of the methylation profile in *ex vivo* bone marrow macrophages and dendritic cells in the H2-Ab1 locus (Target region I). **h.** Experimental procedure for the transduction of bone marrow derived dendritic cells and macrophages with lentiviruses. **i.** Assessment of the transduction efficiency of the lenti-transduced cells at the time point of Day 2 with lenti-mCherry-gRNA viruses through sorting. **j.** mCherry assessment of parameters in transduced cells**. k.** Frequency of total mCherry cells in the bulk transduced *ex vivo* cultures (n=3 per condition). Data is expressed by mean ± SEM. Unpaired t-test was performed in f, j, k. One-way Anova with Tukey’s multiple corrections tests was applied to c-e. Non-significant values are not presented, *, p<0.05. Scale bar: 100um.

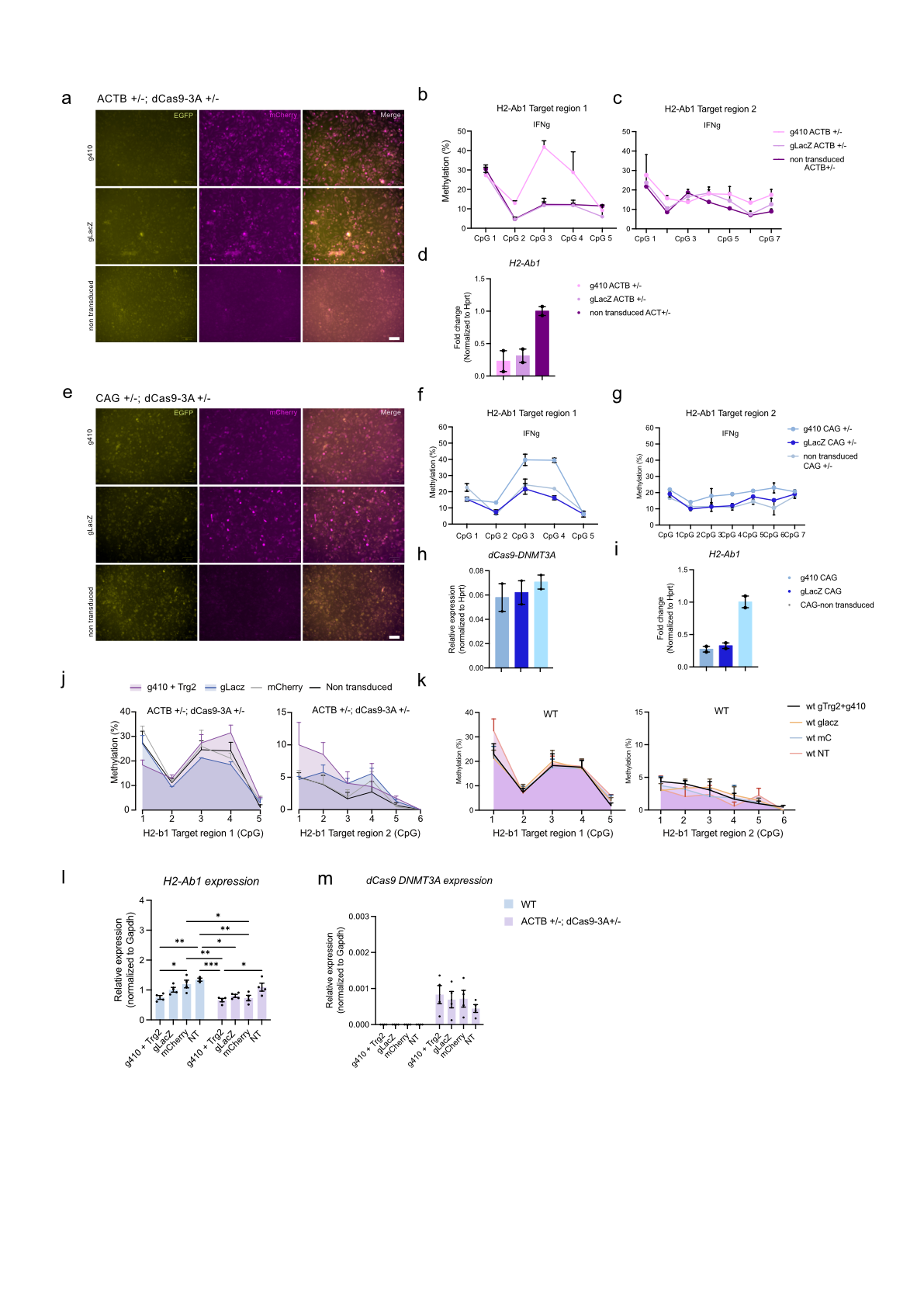
**Supplementary Fig. 15 I Tranduction of *ex vivo* bone marrow-derived macrophages can induce methylation upon mIFNg stimulation**. **a,e.** Fluorescence images from the transduction of ACTB +/-;dCas9-3A +/- and CAG+/-;dCas9-3A+/- cells at day 9 of culture. **b,c,f,g.** Methylation profile of the IFNg stimulated and transduced cells in Target region 1 and 2 (n=2 technical replicates per condition) in ACTB+/- and CAG+/- cells. **d,h,i.** RT-qPCR of the *H2-Ab1* expression with IFNg stimulation h) and *dCas9-DNMT3A* expression of stimulated macrophages (n=2 technical replicates per condition). **j,k.** Methylation profile at the Target region 1 and 2 of the H2-Ab1 locus in ACTB +/-; dCas9-3A +/- and WT transduced and stimulated macrophages upon application of different high titer viruses (n=4 biological replicates per condition). **l,m.** RT-qPCR of *H2-Ab1* and *dCas9-DNMT3A* from the lentivirally transduced cells. Two-way ANOVA with Tukey’s multiple correction testing was performed in l. *, p< 0.05. Scale bar=100 um (a,e).

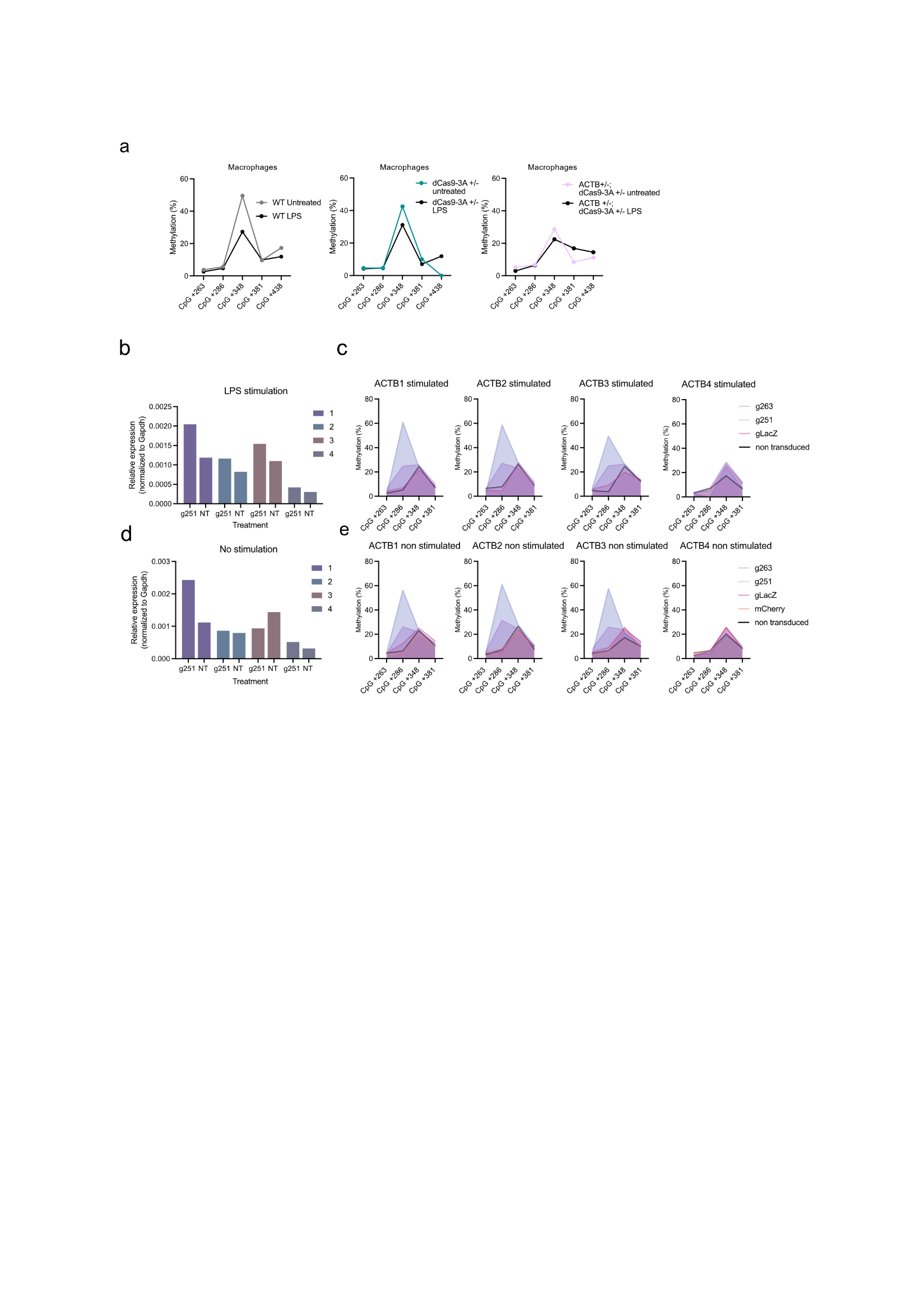

**Supplementary Fig. 16 I** **a**. Assessment of the methylation profile in the IL-6 locus in untreated and 100ng/ml treated with LPS macrophages from ACTB+/- and ACTB-/-; dCas9-3A+/- and WT animals. Individual qPCR analysis of the *dCas9-DNMT3A* expression (**b, d**) and pyrosequencing methylation (**c, e**) in four individual ACTB+/-; dCas9-3A+/- (numbered 1 to 4) animals with or without 10ng/ml of LPS for 6 hours compared to the non-transduced cells (NT).

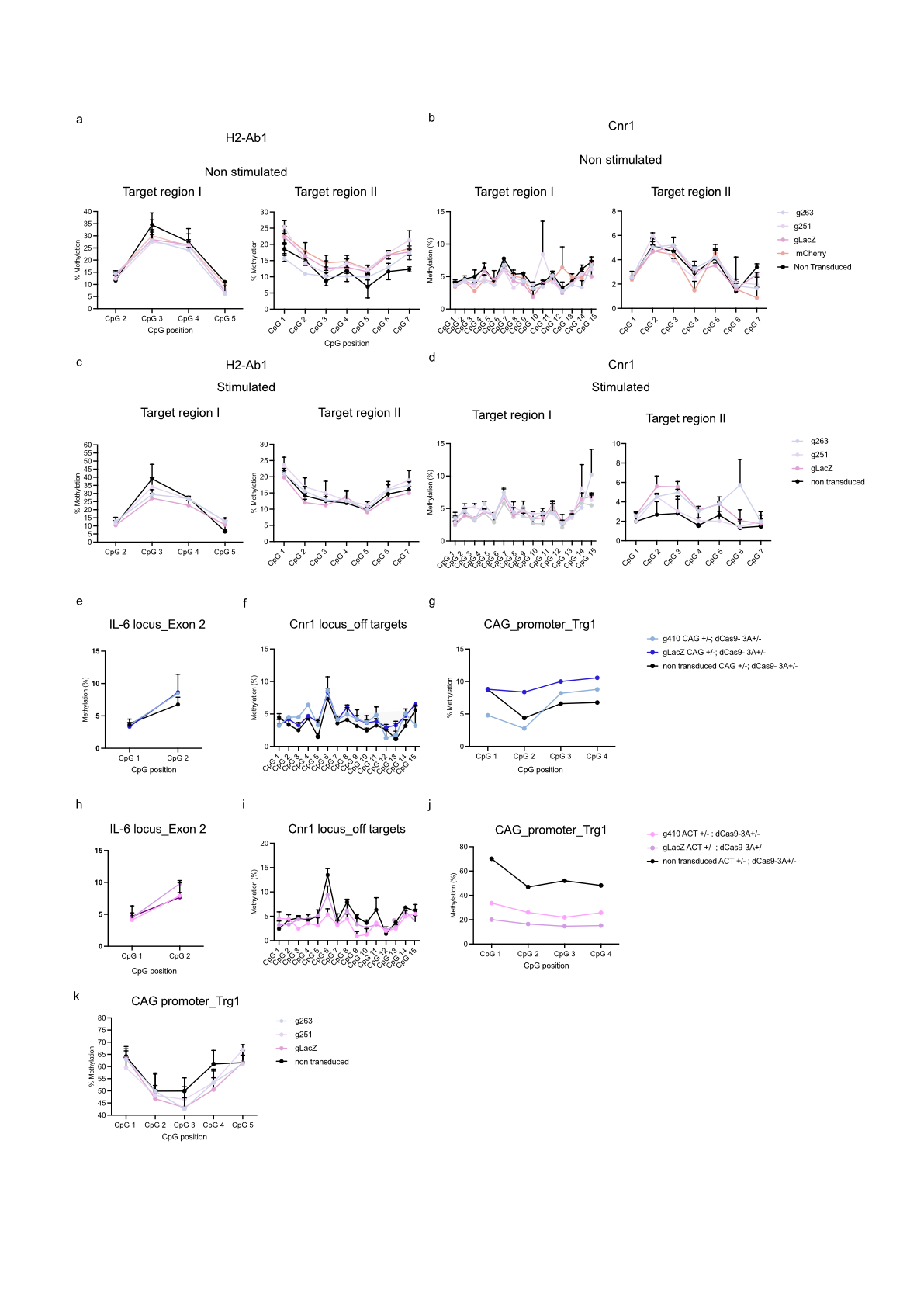

**Supplementary Fig. 17** I **No off-target effects in the lentivirally transduced ex vivo macrophages of the IL-6 locus. a-d.** Methylation analysis by pyrosequencing of *H2-Ab1* (**a, c**) and *Cnr1* (**b,d**) loci in macrophages of ACTB+/-; dCas9-3A+/- animals transduced with Il6-targeting gRNAs (n= 2-3 per condition), unstimulated (a, b) or stimulated with LPS. **e-j**. Methylation analysis by pyrosequencing of *Il6, Cnr1* and *CAG* promoter loci in the non-stimulated macrophages lentivirally transduced with *H2-Ab1*-targeting gRNAs in the CAG+/-; dCas9-3A+/- (**e-g**) and in ACTB+/-; dCas9-3A+/- (**h-j**) animals (n=2 per condition, except j, g n=1 in h.j and n=1 in g). **k**. Methylation profile of CAG promoter in the stimulated lentivirally transduced macrophages of the ACT+/-; dCas9-3A+/- background (n=3 per condition). Data are represented as Mean ± S.E.M. P-values are denoted: *p<0,0332, **p<0.0021, ***p< 0,0002, ****p< 0,0001. Two-way ANOVA with Tukey’s multiple corrections test was performed in (a,b,c,d,k). Non-significant p-values are not represented.

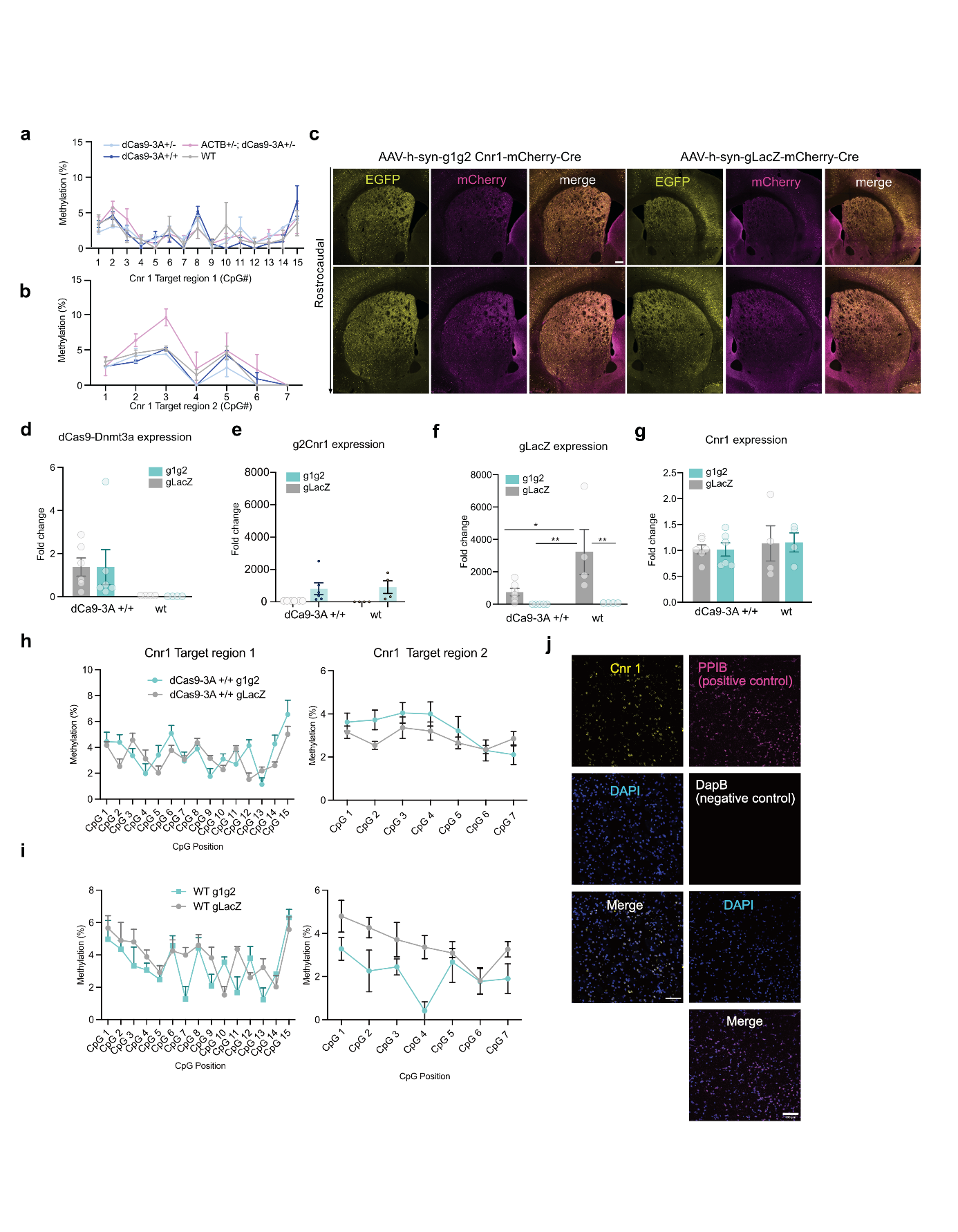

**Supplementary Fig. 18** I **Assessment of the AAV-g1g2-Cnr1-mCherry and AAV-gLacZ-mCherry injections in the brain of different mouse strains.** a,b) Pyrosequencing analysis in the Cnr1 locus (Target region I and II) in the dorsolateral part of the striatum of dCas9-3A +/-, dCas9-3A +/+, ACT+;dCas9-3A +/- and WT animals. c) Representative immunofluorescence images across the dorsolateral striatal axis of an dCas9-3A +/- animals that has received bilateral injections of AAV-h-syn-g1g2 Cnr1-Cre-mCherry and AAV-h-syn-gLacZ-Cre-mCherry, as a control. d) qPCR assessment of the induction of the cassette, dCas9-DNMT3A-EGFP, after intracerebral injections of AAV-g1g2-mCherry-Cre and AAV-gLacZ-mCherry-Cre viruses in the dorsolateral part of the striatum in dCas9-3A +/+ and WT animals n dCas9-3A +/+= 6, n WT= 4 e,f) Assessment of the expression of g2 and gLacZ RNAs in bulk brains of dCas9 -3A+/+ and WT n dCas9-3A +/+= 6, n WT= 4). g) Cnr1 (fold change) in bulk isolated dorsolateral striata from dCas9-3A +/+ and WT animals (n dCas9-3A +/+= 6, n WT= 4) injected with AAV-Cnr1 and AAV-gLacZ viruses. h, i) Pyrosequencing analysis in the Cnr1 locus of the bulk isolated striata in dCas9-3A +/+ and WT animals (n dCas9-3A +/+= 6, n WT= 4). j) RNAscope image for the assessment of the expression of *Cnr1* in the dorsolateral striatum of WT animals. PPIB was used as a positive control and DapB as a negative control for the RNAscope experiments. Data are represented as Mean ± S.E.M. P-values are denoted: *p<0,0332, **p<0.0021, ***p< 0,0002, ****p< 0,0001. Two-way ANOVA with Tukey’s multiple corrections test was performed in (d,e,f). Non-significant p-values are not represented. Scale bar: 200um (a) and 100um (j).
